# Multifaceted roles of extracellular vesicles in *Agrobacterium fabrum* C58 lifestyles

**DOI:** 10.64898/2026.03.19.713014

**Authors:** Timothée Zannis-Peyrot, Fanny Nazaret, Deniz Sarigol, Jeanne Dore, François-Xavier Gillet, Vincent Gaillard, Gilles Comte, Isabelle Kerzaon, Céline Lavire, Ludovic Vial

## Abstract

Bacterial extracellular vesicles (EVs) constitute a key driver of interspecies and inter-kingdom communication, and shape bacterial ecology, yet their role as a dynamic delivery system remains underexplored. Here, we show that the plant pathogen *Agrobacterium fabrum* C58 modulates its EVs in response to virulence-inducing conditions. Our multi-omics analysis revealed that these virulence-state EVs are significantly enriched in effectors from the Type IV secretion system and toxins from the Type VI secretion system, which were previously known to be delivered by conventional contact-dependent mechanisms. We demonstrate that these EVs can directly transfer virulence effectors into plant host cells, enhancing tumor formation. Furthermore, we show that these EVs can interact with and influence the development of several environmental bacteria. Finally, *A. fabrum* C58 EVs elicit distinct plant host metabolome responses compared to whole cells. Our findings establish EVs as a crucial and dynamic component of bacterial virulence and inter-kingdom communication, providing a new perspective on how bacteria adapt to and manipulate their environment.

## Introduction

The environment constantly shapes bacteria behaviors, ecology, and evolution, and in turn modulates interaction outcomes at community and host interfaces. In response to host or environmental cues, bacteria acquired lifestyle flexibility, allowing them to colonize specific habitats^1,2^. This flexibility is particularly relevant in the plant environment, where bacteria are required to adapt to conditions imposed by the host and the associated microbiota^3^. Indeed, some plant-interacting bacteria known to adapt to various ecological niches could be used as suitable models to study how life-specific conditions shape bacteria physiology. Bacteria from the genus *Agrobacterium* can be commensal and are highly abundant in the soil and the rhizosphere^4^. However, it is only when it harbors the Tumor-inducing plasmid (pTi) and in response to wound-associated host-like signals, that it switches to a pathogenic lifestyle associated with changes in bacteria behavior, physiology and metabolism^5,6^. Indeed, virulence-inducing conditions drive the transfer of pTi-localized T-DNA fragment and Vir proteins/effectors from bacteria to plant cells through a Type IV secretion system (T4SS), leading to crown-gall tumor formation that constitute a novel host-derived niche colonized by adapted bacteria^7^. Virulence-inducing conditions also induce the expression of *A. fabrum* Type VI secretion system (T6SS) that is used to compete against adversaries both *in vitro* and *in planta*, mediating contact-dependent killing that may influence plant-associated community composition^8,9^. These findings place *Agrobacterium*, and more specifically *A. fabrum* C58, as a model for studying bacterial environment-driven adaptation, linking lifestyle switches to secretion- and competition-based strategies.

Such strategies allow bacteria to suitably interact and respond to their environment, and generally involve the production and release of specific molecules in the bacterial surroundings. In addition to soluble factors, bacteria are able to produce extracellular vesicles (EVs), characterized as lipid nanospheres (∼50–400 nm) derived from bacterial membranes^10^. By protecting the encapsulated cargo from environmental constraints, they enable long-distance transport and delivery of a concentrated payload usually composed of proteins, lipids, nucleic acids and metabolites^11^. Once released into the environment or internalized by recipient cells, this heterogeneous cargo can modulate gene expression, signaling, metabolism and community dynamics, thereby shaping intra- or inter-species interactions^12–14^. Bacterial EVs are mostly studied in the context of animal-bacteria interactions and have been shown to play dual roles in both bacterial pathogenicity and host immune response elicitation^14,15^. More recently, EVs from plant-interacting bacteria gained increasing importance as they were shown to also play both roles in plant-bacteria interactions, making them new key actors of phytobacterial ecology^16,17^. However, the roles of EVs from plant-interacting bacteria under environmental conditions are still poorly explored.

Given the importance of plant environment in shaping both *A. fabrum* C58 lifestyle and EVs properties in several phytobacteria models, we asked how host-associated contexts would modify *A. fabrum* C58 EVs and, in turn, the produced EVs would shape *A. fabrum* C58 interactions with the plant or its microbial partners. To test these hypotheses, we first characterized the influence of virulence-inducing conditions on *A. fabrum* C58 EVs content and compared it to whole-cell lysates (WCL) by performing multi-omics LC-MS² analyses. Thereafter, we assessed the effect of EVs produced in minimal or virulence-inducing conditions on environmental bacteria development using both bacterial challenge and binding assays. On the plant side, we show that virulence factor from bacterial EVs can be transferred to plant cells. We used the plant model *Solanum lycopersicum* to monitor the impact of EVs from each condition on bacterial virulence and crown-gall formation. We mapped host metabolome responses to EVs from both conditions by untargeted LC–MS² analyses, comparing these with responses elicited by whole living bacterial cells.

## Methods

### Bacterial strains and growth condition

*A. fabrum* C58 wild-type (WT) was cultivated in YPG rich medium (Yeast extract 5 g.L⁻¹; bacterial peptone 5 g.L⁻¹; glucose 10 g.L⁻¹) or AB minimal medium^18^ pH 7.0 supplemented with glucose (10 mM) as a carbon source (**AB Glucose**), under agitation (180 rpm) when liquid medium was used, or on agar plates (agar 15 g.L⁻¹). To induce virulence, *A. fabrum* C58 was cultivated in AB Glucose medium supplemented with acetosyringone (15 µM), and with a modified pH at 5.5 (**AB Virulence**) (**Text S1.**). For the rest of the experiments, cells and EVs will be named **Glu** when originating from **AB Glucose** medium, and **Vir** when originating from **AB Virulence** medium. 1A medium was used to select *A. fabrum* C58 after in planta inoculations^19^. Bacterial strains (**Table S1a**) used in bacterial challenge were cultivated in Trypton Soy Broth (TSB) or TSB 1/10 strength under agitation (180 rpm) when liquid medium was used, or on agar plates (agar 15 g.L⁻¹).

### *A. fabrum* C58 virulence induction assay

Virulence induction was verified using an *A. fabrum* C58 reporter strain carrying the pOT1e-*virB-Egfp* plasmid^6^ grown in AB Virulence medium and compared to cells grown in AB Glucose medium. Cells were inoculated (OD600nm = 0.025) in 1 L of medium up to 31 h at 28°C under agitation. Every hour between 24 and 31 h of culture, 600 µL of culture was sampled and distributed into three wells of a 96-well microplate, each containing 200 µL, for fluorescence reading using TECAN plate reader (ex: 485 nm; em: 530 nm).

### Plasmids and strains construction used for the split-GFP system

The VirE2-sfGFP11 fusion protein was constructed using Gibson Assembly cloning in a pJQ200sk suicide-plasmid, based to the method of **Li *et al.,* 2014**. Briefly, the sfGFP11-coding sequence was inserted into *virE2* at Pro54 permissive site^20^ using overlapping PCR primers. The resulting DNA fragments were inserted into the pJQ200sk *sacB*-suicide plasmid^21^ to generate pJQ200sk-VirE2-sfGFP11. In parallel, sfGFP1-10 coding sequence was amplified from pET22b-sfGFP1-1022 using overlapping primers and cloned by Gibson Assembly into the broad-host-range plasmid pBBR1-MCS5 under the control of a T7 promoter. All constructs were confirmed by PCR and sequencing analysis. Plasmids were introduced into *A. fabrum* C58 WT by electroporation. Single recombinants were selected on YPG media containing 20 μg.mL^-1^ gentamicin, and finally double recombination events were identified by sucrose resistance on YPG media supplemented with 5% sucrose. The pBBR1-sfGFP1-10 plasmid was introduced into the *A. fabrum* C58 WT and the VirE2-sfGFP11 expressing strain, and transformants were verified by PCR. All plasmids and primers used in this study are listed in **Table S1b**. To control the assembling ability of sfGFP1-10 and VirE2-sfGFP11 proteins, *A. fabrum* C58 WT carrying pBBR1-sfGFP1-10 (as a negative control) and the VirE2-sfGFP11 strain carrying pBBR1-sfGFP1-10 were grown in AB Virulence conditions, and fluorescence was measured in a TECAN plate reader (ex: 485 nm; em: 530 nm).

### EVs isolation by tangential flow filtration and gradient density ultracentrifugation

Bacterial cells grown for 31 h to an OD600nm of 0.8 and 0.4, in AB Glucose and AB Virulence respectively. Cells were pelleted from 2 L of culture in a Beckman Avanti J-E series centrifuge (JLA-14 rotor; 12,000 x *g*; 10min). The supernatant (2 L) was collected and filtered (0.45 µm RC, Millipore Durapore) to remove any residual bacterial cells. EVs were then concentrated to 30 mL using tangential flow with a 100 kDa cut-off filter (Sartorius). EVs were pelleted from the concentrate in a Beckman Optima LE-80K ultracentrifuge (70Ti rotor; 200,000 x *g*; 2h30; 4°C). The EVs pellet was resuspended in 20 mL of sterile dH2O. When needed, EVs were labeled with FM4-64 (1 µg.mL⁻¹) 30 min at 28°C in the dark. Concentrated EVs samples were then layered on top of an Optiprep gradient and centrifuged (50%, 40%, 30%, 20%; 1 mL each; SW32 rotor; 150,000 x *g*; 18 h; 4°C). Fractions were collected from the top of the ultracentrifuge tube, washed in 30 mL of sterile dH2O and centrifuged (70Ti rotor; 200,000 x *g*; 4 h; 4°C). The supernatant was discarded, and pellets were resuspended in 1 mL sterile dH2O before protein and lipid quantification. Samples were plated onto LB agar medium to verify the absence of contamination after each EVs purification. EVs samples were used immediately for content analysis and plant experiments. Lipid content was assayed using FM4-64 microplate assay. Ten microliters of EVs samples were mixed with 200 µL of FM4-64 (1 µg.mL⁻¹) for 30 min at 28°C in the dark. Fluorescence was measured in a TECAN plate reader (ex: 515 nm; em: 635 nm) and compared to a standard liposome curve prepared as described previously^23^.

### Electron Microscopy

For scanning electron microscopy, 2 mL of cells isolated from cell cultures (used for EVs extraction) were centrifuged (600 × *g*; 3 min). The supernatant was discarded, and cells were resuspended in 50 μL of Phosphate Buffer (PB) 0.1 M pH 7.4. Samples were fixed for 3h in PB containing 2% glutaraldehyde (*v/v*) and then filtered through a 0.2 μm filter. Samples were washed 3 times for 20 min in PB 0.2 M prior to post-fixation in 1% osmium tetroxide (*v/v*) (diluted in PB 0.1 M) for 45 min. Samples were rinsed several times with water, then dried overnight in 20% acetone (*w/v*) in a vacuum bell. Samples were incubated in pure acetone for 1 h at room temperature before being dried with Critical Point Dryer Leica EM CPD300 (= V/med/120s/5/12/med/slow 100%) for 2 h. Preparations were mounted on stubs and sputter-coated with carbon and then analysed in a scanning electron microscope model FEG Merlin COMPACT. For transmission electron microscopy (TEM), 10 µL of GDUC-purified EVs samples were dropped on a hydrophilized copper Formvar grid (400 mesh) for 2 min. Fixed EVs were then contrasted with 1% tungsten disilicide (*w/v*) for 1 min before the grids were washed for 1 min with H2O and left to dry. Samples were analysed using the transmission electron microscope model JEM −1400 Flash.

### Nanoparticle Tracking Analysis

Nanoparticle Tracking Analysis (NTA, Zetaview, ParticleMetrix) was used to estimate the EVs concentration and size distribution in purified samples. DiO labeled (1 µg.mL⁻¹) EVs samples were diluted 1000 times in dH2O and 1 mL of each resulting dilution was injected into the NTA (ZetaView). The ZetaView software (8.05.16 SP3 version) was used to acquire and process each sample video. The instrument was set at a temperature 25°C, sensitivity 80, 488 nm laser wavelength. For DiO detection, the instrument was run in fluorescence mode with 488 nm excitation and a 500 nm long-pass emission filter, restricting tracking to DiO-positive particles. AB Glucose medium and dH2O were used as negative controls. The amount of EVs in the sample was calculated using the correspondence between EVs particle numbers calculated by NTA and total lipid amount quantified by FM4-64 assay.

### Bacteria and EVs content analysis

Three biological replicates of *A. fabrum* C58 cultivated in Glucose or Virulence (WCLGlu and WCLVir) condition and their respectively produced EVs (EVGlu and EVVir) were subjected to Metabolite, Protein and Lipid Extraction (MPLEx) protocol^24^. The detailed methodology for protein, polar and apolar metabolites extraction and analysis is described in **Text S2-4**. Following MPLEx protocol, dried protein fractions were processed following single-pot, solid-phase-enhanced sample-preparation (SP3) protocol^25^. Peptide concentration was determined by quantitative fluorometric peptide assay in order to inject 200 ng of peptides into the nanoLC-MS² system. Digested samples were analyzed by label-free quantitative approach (LFQ) on the Exploris 480 mass spectrometer coupled to a Vanquish NEO nanoLC system (Thermo Scientific). Raw data were processed with the Proteome Discoverer 3.1 software through the CHYMERIS 3.0 search engine against the *Agrobacterium fabrum* database (Uniprot, January 2025). Protein quantitation was performed with precursor ions quantifier node in Proteome Discoverer 3.1. Dried polar and apolar fractions were solubilized at 10 mg.mL^-1^ in MeOH 50:50 (*w/v*) and pure MeOH respectively. Metabolite analyses were performed by UHPLC-DAD-ESI-MS/MS-QTOF on an UHPLC Agilent 1290 coupled to a UV-vis DAD and a high-resolution Q-TOF 6546 mass spectrometer (Agilent Technologies, Santa Clara, CA, USA). The device was managed by the Agilent MassHunter DataAcquisition 11.0. Data was processed using MzMine 4.5.0^26^.

### Bacterial challenge and EVs uptake assays

Bacterial strains (**Table S1a**) were grown overnight in 10 mL of TSB 1/10 at 28°C. Overnight cultures were centrifugated (7000 x *g*; 10 min) and washed to OD600nm = 1 in Phosphate Buffer Saline (PBS). For bacterial challenge, each bacterial suspension was diluted in 200 µL of TSB 1/10 to OD600nm = 0.025 and inoculated with either 5.10^8^ EVs particles of EVGlu or EVVir. Cell suspensions inoculated with sterile dH2O were used as negative controls. Bacterial growth (OD600nm) was monitored in triplicate in Bioscreen plates using Bioscreen plate reader (3 days, 28°C, shaking every 20 min before measurement). The experiment was done in n = 2 biological replicates. For EVs uptake assay, 600 µL of each bacteria suspension (OD600nm = 1) were inoculated with either 5.10^8^ EVs particles of EVGlu or EVVir labeled with FM4-64 (1 mg.mL^-1^) in LoBinding 2 mL Eppendorf tubes at 28°C for 1 h in the dark without agitation. Cell suspensions inoculated with sterile dH2O and PBS alone were used as negative controls. After 1 h, suspensions were washed twice with PBS (7000 x *g*; 10 min) to remove non-binded EVs. Residual fluorescence was measured in triplicate in a 96-well microplate using a TECAN plate reader (ex: 515 nm; em: 635 nm). The experiment was done in n = 3 biological replicates.

### Arabidopsis thaliana growth conditions and inoculation

Seeds of *A. thaliana* transgenic lines expressing CYTO-sfGFP1-10 (pUBQ10:sfGFP1-10OPT:NOSt) obtained from the Nottingham Arabidopsis Stock Centre (NASC ID: N69831) were surface-disinfected and germinated *in vitro* in Murashige and Skoog (MS) ½ medium (Sigma Aldrich). After stratification during 2 days at 4 °C in the dark, seeds were germinated and grown under a 16 h light / 8 h dark photoperiod in a growth chamber at 22 °C. Finally, 1-week old transgenic *A. thaliana* plants were transferred into a fresh MS ½ medium. Half of the root individuals were carefully wounded with a sterile syringe two weeks after germination, while the other half were kept unaffected. Then 1.10^8^ EVs particles purified from either *A. fabrum* C58 WT or *A. fabrum* C58 VirE2-sfGFP11 grown in AB Virulence medium, or sterile dH2O were inoculated. Plants were placed back in the growth chamber for 24 h before imaging.

### Confocal microscopy and image analysis

Zeiss LSM 800 microscope was used to visualize sfGFP and FM4-64 fluorescence (excitation: 488 nm; emission collection: 492–600 nm for sfGFP; excitation: 506 nm; emission collection: 600–700 nm for FM4-64) and transmission light on 24 hpi inoculated *A. thaliana* roots with bacteria or purified EV. Images were captured using EC Plan-Neofluar 10x/0.30, Epiplan 20x/0.40 or Plan-Apochromat 40x/1.3 Oil DIC (UV) VIS-IR M27 objectives, and detector gains fixed at 784 V for sfGFP and 502 V for FM4-64. Fluorescence of sfGFP and FM4-64 was acquired in Zstack acquisition mode on ZEN 3.12 software (© Carl Zeiss Microscopy GmbH 2002-2011 – OME). ImageJ (Fiji) with the Bioformat plugin was used to process all the images in .czi. Each *A. thaliana* imaging experiment included n = 4 roots per condition, and the experiments were done three times independently. At least 4 images per different plant root were used to evaluate fluorescence intensity differences between conditions.

### Tumor colonization assay

Three weeks old *Solanum lycopersicum* var. Marmande grown in greenhouse were wounded by incising stems on 2 cm. Wounds were inoculated with 5 µL of dH2O containing either 1.10^8^ EVs from the EVGlu or EVVir conditions, or 5.10^4^ *A. fabrum* C58 cells mixed with 1.10^8^ EVs particles from the EVGlu or EVVir conditions (n = 9). Plants inoculated with dH2O, *A. fabrum* C58 cells alone and ultracentrifugation supernatant were used as control. Plants were randomized and left to grow in the greenhouse (16h day/8h night cycle). After 3 weeks, tumors were harvested and immediately used for tumor count, *A. fabrum* C58 colonization quantification and nopaline quantification. The experiment was independently repeated three times over 3 different weeks. After the tumor count, the stem tomato samples were cut and half freeze-dried for nopaline quantification while the other half was used for bacterial colonization quantification. To determine bacterial colonization levels, samples were grounded and serial plated onto 1A medium using a spiral plater (2 plates per sample, 3 dilutions per plate) (EasySpiral; Interscience, Saint-Nom-la-Bretèche, France). After 48 h of incubation at 28°C, colonies were counted using a Scan1200 camera coupled with a computer system for CFU determination. CFU numbers were normalized by the dry mass weight of the sample and log10 transformed. The detailed methodology for targeted nopaline extraction is provided in **Text S5**. Briefly, dried samples were crushed and subjected to two successive extractions with 20% MeOH. Extracts were then solubilized at 10 mg.mL^-1^ in MS-grade H2O. Nopaline content was analyzed by UHPLC-ESI-MS QTOF on a UHPLC Agilent 1290 coupled to a DAD and a high-resolution Q-TOF 6530 mass spectrometer (Agilent Technologies, Santa Clara, CA, USA). The device was managed by Agilent Mass Hunter Data Acquisition version 11.0. These analyses, performed to detect and quantify nopaline in the stem/tumour extracts, followed the exact protocol and methodology described previously^27^.

### *S. lycopersicum* amino acids and metabolites analysis

Seeds of *S. lycopersicum* var. Marmande were disinfected and grown as described previously^28^. After 14 days of growth, roots were inoculated with either sterile water supplemented with iodixanol 0.01% (Control) or 1.10^7^ *A. fabum* C58 cells cultivated in Glucose or Virulence condition (C58Glu and C58Vir respectively) or 5.10^8^ of EVs particles (EVGlu and EVVir). Sixty plants per condition were inoculated and put back in the growth chamber. Before plant harvest, an aliquot of the plant growth medium was plated to assess the sterility of the experimental conditions. Eighteen hours post-inoculation, plants of each condition were randomly gathered into six pools of ten plants each (to reduce variability and to ensure sufficient materials). Roots and aerial parts were harvested separately before being quickly frozen in liquid nitrogen, freeze-dried and crushed. Samples were subjected to successive methanolic extractions (see details in **Text S6**). Extracts were then solubilized at 10 mg.mL^-1^ in MeOH/H2O 80:20 (*v*/*v*). The plant free amino acids were analyzed by HPLC-UV/DAD-FLD on an HPLC Agilent 1100 coupled to a DAD and a fluorometric detector (FLD) as described previously^29^. The device was managed by Agilent ChemStation LC Rev. B.04.03-SP2, and the data was processed with Mass Hunter Qualitative Analysis B.07.00. The analysis of tomato specialized metabolites was performed by UHPLC-ESI-MS QTOF on a UHPLC Agilent 1290 coupled to a DAD and a high-resolution Q-TOF 6546 mass spectrometer (Agilent Technologies, Santa Clara, CA, USA). The device was managed by Agilent MassHunter Data Acquisition version 11.0. Data was processed using MzMine v4.5.0 for specialized metabolites profiles matrix obtention^26^. The detailed methodology for *S. lycopersicum* amino acids and metabolites content analyses are provided in **Text S7-8**.

### Metabolite annotation and molecular networking

The detailed methodology for *S. lycopersicum* metabolites annotation and molecular network clustering are provided in **Text S9**. EVs and plant metabolites annotation was carried out using SIRIUS software version 6.1^30^. Metabolite classes and synthesis pathways were determined by the CANOPUS module^31^. Metabolite identification was either carried out using the CSI:FingerID module^32^ or manually searched in the in-house SeMetI database and/or in the public databases Lotus, Knapsack, GNPS, CasSciFinder and the literature. To facilitate and deepen the identification of plant metabolites, a molecular network was constructed using the t-SNE algorithm in the MetGem software v1.5.2^33^.

### Statistical analysis

All statistical analyses were conducted using R software (v4.4) and the plots were created using the ggplot2 package (v3.5.1), unless otherwise stated. Final figures were designed using InkScape (v1.4). Detailed statistical analyses are provided in **Text S10**.

### EV Tracker metrics

We have submitted all relevant data of our experiments to the EV-TRACK knowledgebase^34^, scoring an EV-metric score of 91%. You may access and check the submission of experimental parameters to the EV-TRACK knowledgebase via the following URL: http://evtrack.org/review.php. Please use the EV-TRACK ID (EV250069) and the last name of the first author (Zannis-Peyrot) to access our submission.

### Data availability

The mass spectrometry metabolomics data have been deposited to the Center for Computational Mass Spectrometry repository (University of California, San Diego) via the MassIVE tool with the dataset identifier MassIVE MSV000099970. EVs and bacteria metabolites and lipids data are available at https://www.ebi.ac.uk/metabolights/MTBLS13797. Plant metabolites study and data are available at https://www.ebi.ac.uk/metabolights/MTBLS13796.

## Results

### Virulence-inducing conditions do not influence EVs production

To compare the effect of dual *A. fabrum* C58 lifestyles on EVs production and cargo, EVs were purified from a growth medium mimicking the conditions needed for *A. fabrum* C58 virulence and compared to a classic medium with a carbon source usually found in plant environment (glucose). Growth duration of 31 h was chosen as a sampling time as it is in the late-exponential phase window for both growth conditions and showed a statistical increase in virulence induction of *A. fabrum* C58 in AB Virulence compared to AB Glucose condition (**Fig. S1**). Visualization of *A. fabrum* C58 cells from growth medium by scanning electron microscopy revealed the presence of EVs of a size approximately 100 nm and some blebbing from the outer-membrane of cells (**Fig. 1a-b, Fig. S2**). To help EVs purification, EVs pellets obtained after the first ultracentrifugation round were labeled with DiO lipophilic dye (1 mg.mL^-1^) (**Fig. S3**). Nanoparticle tracking analysis (NTA) revealed that *A. fabrum* C58 produced EVs of ∼124 nm and ∼110 nm in size in AB Glucose and AB Virulence respectively, as confirmed by TEM analysis, at concentrations of 7.7.10^9^ particle.mL^-1^ in both conditions (**Fig. 1c-f**). No particles were detected in the other gradient layers or AB Glucose medium. Finally, zeta-potential (ZP) measurement indicated that purified EVs from *A. fabrum* C58 had a relatively low ZP (−67.1 mV in AB Glucose and -55.7 mV in AB Virulence), assessing the stability of purified EVs samples. Overall, statistical analysis of EVs size, concentration and ZP between AB Glucose and AB Virulence conditions didn’t reveal any significant difference, indicating that virulence-inducing conditions did not influence EVs production and morphology in *A. fabrum* C58.

**Fig. 1:**
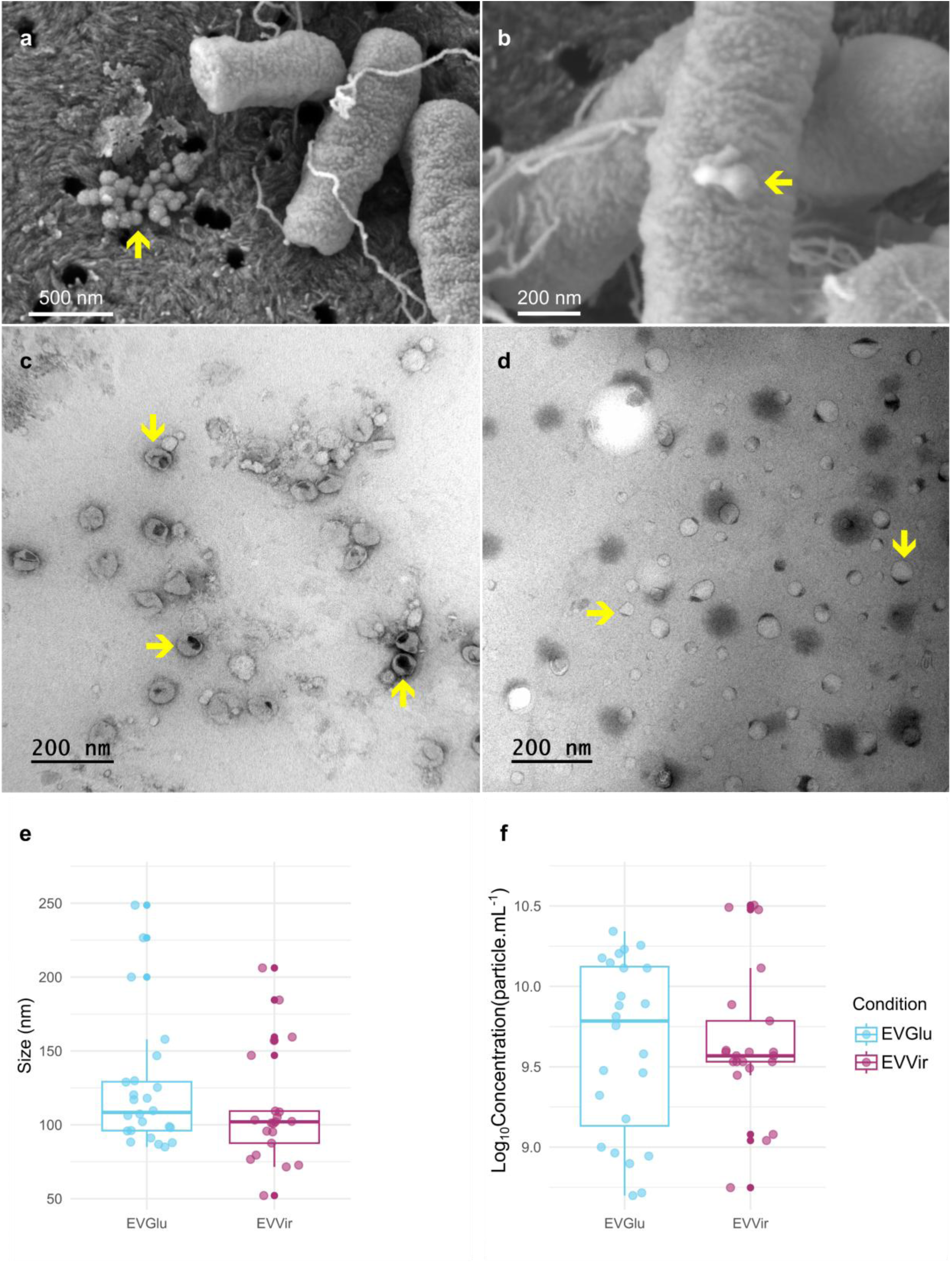
Production of EVs by *A. fabrum* C58 in minimal and virulence-inducing conditions. (**a**) Scanning electron micrographs of *A. fabrum* C58 cells and its EVs. Scale bar 500 nm. (**b**) Close up of a scanning electron micrograph displaying an EV blebbing from the outer-membrane of the bacterial cell. Scale bar 200 nm. Transmission electron micrographs (TEM) of EVs from *A. fabrum* C58 purified from either (**c**) AB Glucose or (**d**) AB Virulence growth conditions. Yellow arrows indicate the presence of EVs. Scale bars 200 nm. Comparison of (**e**) EVs size (nm) and (**f**) EVs concentration (particle.mL^-1^) measured by NTA in purified EVs samples from the AB Glucose or AB Virulence condition. Box plots show median (center line), interquartile range (box), and 1.5× interquartile range (whiskers) and technical replicates from *n* = 3 biological replicates shown as points. Two-sided Student’s *t*-test was used for statistical analysis. No statistical differences between the two groups were observed.

### *A. fabrum* C58 EVs are enriched in outer-membrane proteins compared to whole cell lysates

To define the cargo of *A. fabrum* C58 EVs, Whole Cell Lysates (WCL) and purified EVs were subjected to MPLEx separation protocol and the resulting 3 phases corresponding to proteins, apolar metabolites and polar metabolites were analyzed separately using LC-MS^2^. Out of the 5,344 proteins potentially encoded by *A. fabrum* C58 genome, 2,312 were detected in *A. fabrum* C58 WCL in AB Glucose growth medium (**Table S2c**). Among them, 1,627 (70.37%) were found in EVs samples purified from the Glucose condition (EVGlu) (**Table S2a**). Top 10 GO terms analysis revealed proteins associated to Biological Process (BP) terms such as “translation”, “proteolysis”, “peptide transport”, “cell-wall organization”; Cellular Component (CC) categories corresponding to “cytoplasmic”, “membrane” and “periplasmic space”; and to Molecular Functions (MF) terms including “ATP binding” and “metal ion binding” (**Fig. 2a**). KEGG pathway annotation revealed that most proteins belonged to the KEGG Level B terms “Carbohydrate metabolism”, “Amino acid metabolism” and “Membrane transport” (**Fig. 2b**). Among the most abundant proteins found in EVGlu samples were the OMVs marker proteins Atu0682 (GroEL), Atu2722 (OmpA) and Atu1381 (Omp1), indicating that our EV samples contain Outer-Membrane Vesicles. This was supported by the presence of the outer-membrane protein Atu8019, an *A. fabrum* C58 EVs protein described previously^35^. Consistent with these observations, protein localization prediction revealed a significant enrichment in outer-membrane, extracellular and periplasmic proteins and a significant depletion in both cytoplasmic and inner-membrane proteins in EVGlu samples compared to WCLGlu (**Fig. 2c**). The Microbe-Associated Molecular Patterns (MAMPs) FlaA, FlaB and Ef-TU were also found in EVs samples as it was already described in many studies regarding phytobacterial EVs^16^. LC-MS² analysis of the apolar fraction allowed the detection of 748 ions from apolar compounds in WCLGlu and 616 in EVGlu samples. In EVGlu, ions were mostly assigned to the proposed biosynthesis pathways “Fatty acids”, “Terpenoids” and “Alkaloids”, and mapped to the proposed chemical classes including “Diacylglycerols”, “Glycerophosphocholines” (PC), “Glycerophosphoethanolamines” (PE), “N-acyl amines” and “Triacylglycerols” (**Fig. 2c**, **Table S3a**). Finally, LC-MS² analysis of the polar fraction allowed the detection of 89 ions in WCLGlu and 31 ions in EVGlu assigned to the proposed biosynthesis pathways of “Fatty acids”, “Amino acids” and “Alkaloids” (**Fig. 2e**, **Table S4a-S4c**). Altogether, these results show that *A. fabrum* C58 produces EVs, from which OMVs, carrying a protein and lipid cargo comparable to previously described plant-associated bacteria^16^.

**Fig. 2:**
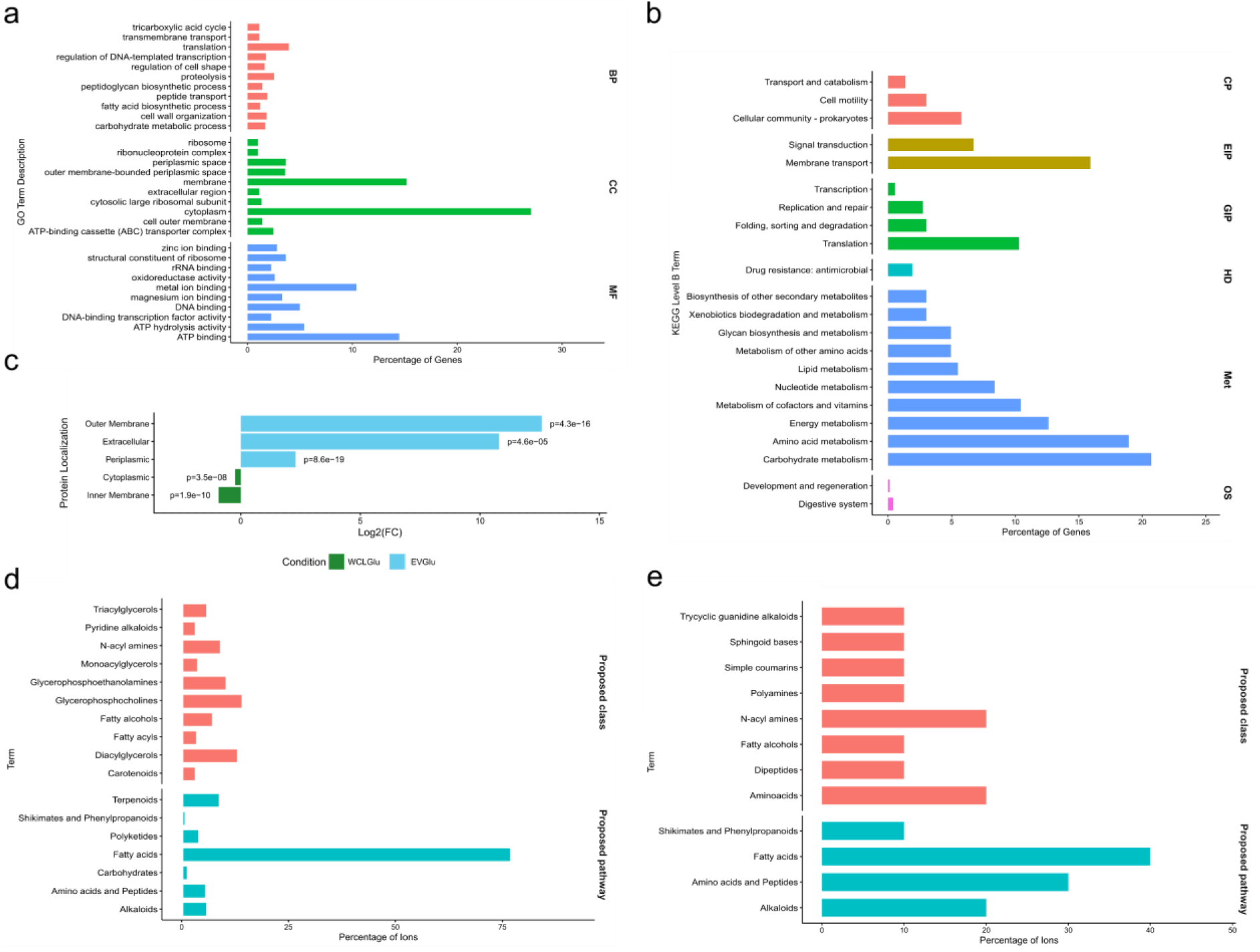
Description of the molecular content of *A. fabrum* C58 EVs. Protein, apolar and polar metabolites content was separated following MPLEx protocol on 3 biological replicates. (**a**) Top 10 GO Terms from the Biological Process (BP; red), Cellular Components (CC, green) and Molecular Functions (MF, blue) categories assigned to the 1627 proteins found in EVGlu protein samples. (**b**) Top 10 KEGG Level B Terms assigned to the KEGG Level A categories Cellular Processes (red), Environmental Information Processes (brown), Genetic Information Processing (green), Human Diseases (cyan), Metabolism (blue) and Organismal Systems (pink) assigned to the 1627 proteins found in EVGlu protein samples. Bars represent the percentage of proteins assigned to each category. (**c**) Comparison of the protein predicted localization between EVGlu and WCLGlu. Protein localization was predicted using DeepLocPro 1.0. Enrichment was calculated using Fisher’s test between the two compared conditions. Protein localizations were considered enriched or depleted when *p* < 0.05. (**d**) Top 10 Proposed Class (red) and Proposed pathway (cyan) of the 616 ions detected in EVGlu apolar metabolites samples. Bars represent the percentage of ions assigned to each category. (**e**) Top 10 Proposed Class (red) and Proposed pathway (cyan) of the 31 ions detected in EVGlu polar metabolites samples. Bars represent the percentage of ions assigned to each category.

### *A. fabrum* C58 EVs are enriched in Type IV secretion–associated proteins compared to the bacteria

While few studies compared the EVs cargoes to the WCL of the bacteria they originate, doing so might reveal unique enriched metabolic pathways and help decipher the ecological roles of EVs. Comparing the content of *A. fabrum* C58 WCL and EVs from the Glucose condition revealed an enrichment in 105 proteins and a depletion of 35 proteins in EVs compared to WCL (**Table S2, Fig. S5a**). GO terms enrichment analysis revealed an enrichment in proteins belonging to the BP “protein secretion by the type IV secretion system” and CC “cell outer membrane” and “type IV secretion system complex”. Out of the 11 structural proteins of *A. fabrum* C58 T4SS, the proteins anchored to the inner membrane Atu6170 (VirB4) and Atu6171 (VirB5), as well as the proteins anchored in the outer-membrane Atu6173 (VirB7), Atu6174 (VirB8) and Atu6175 (VirB9) were found enriched in EVs compared to the WCL (**Fig. 3b**). However, no Type IV secretion system (T4SS) effectors were found enriched in EV samples. The EV marker outer membrane proteins Atu1131 (RopB) and Atu2159 (Omp) were also found enriched in EV Glu samples.

**Fig. 3:**
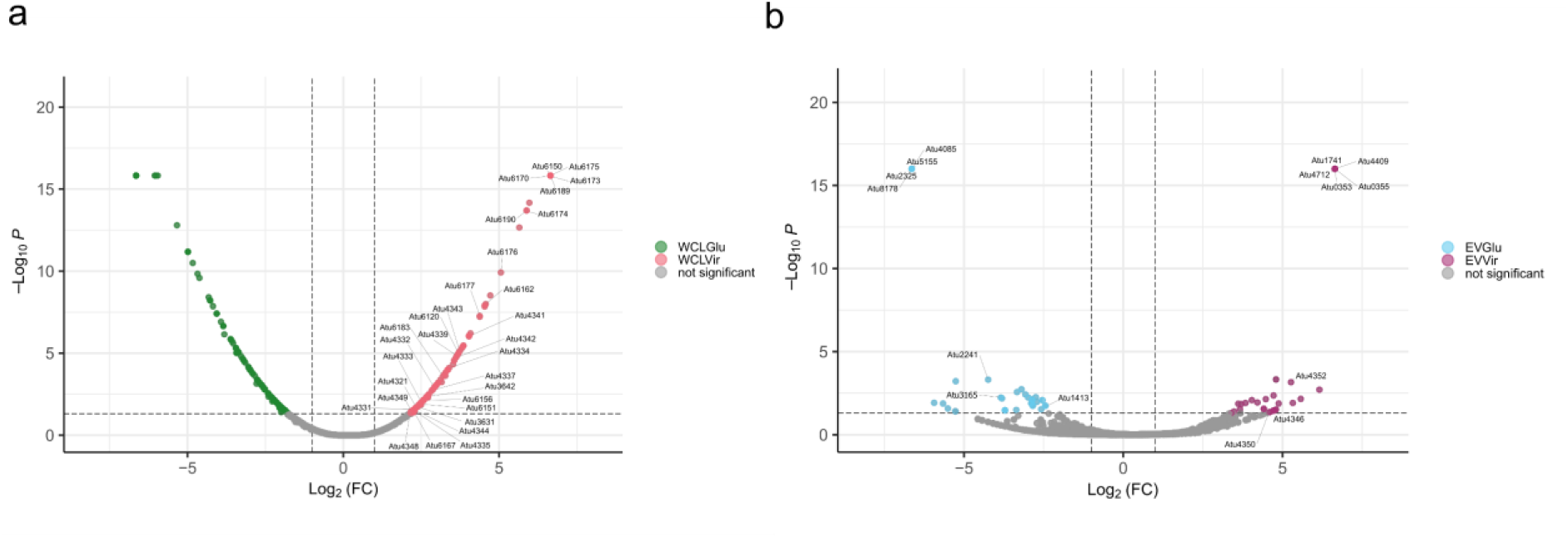
Virulence conditions modulate the protein profiles of *A. fabrum* C58 and its EVs. Volcano plot displaying differential abundance of protein between (**a**) WCLVir (right, red) and WCLGlu (left, green) or (**b**) EVVir (right, purple) and EVGlu (left, cyan) as determined by a Student’s *t*-test with BH correction (*p* < 0.05, Log2(FC) > |0.5|). Data represent the mean value of *n* = 3 biological replicates.

### Virulence medium changes the cargo of *A. fabrum* C58 EVs

As virulence induction modifies the lifestyle of *A. fabrum*, we wanted to investigate its effect on produced EVs cargoes. Comparison between WCLGlu and WCLVir analysis revealed an enrichment in many proteins encoded by the pTi-located *vir* regulon and the T6SS regulons and confirmed virulence induction. These results are in accordance with previous proteomic and transcriptomic studies^36,37^. EVVir samples shared most of its highly abundant proteins with EVGlu samples, especially EVs marker proteins like Atu0682 (GroEL) and Atu1381 (Omp1) (**Table S2b**). However, virulence-inducing conditions induced the significant enrichment of 28 proteins and the depletion of 26 proteins in *A. fabrum* C58 EVs compared to the Glucose condition (**Table S2e**, **Fig. 3b**). Three secreted proteins belonging to *hcp* operon from the T6SS of *A. fabrum* C58 were found significantly enriched in EVVir samples. Especially, the DNase Atu4350 (Tde1) and the antitoxin Atu4346 (Tai) were found enriched. Their respectively associated proteins Atu4351 (Tdi1) and Atu4347 (Tae) were not found significantly enriched in EVVir despite being present in high abundance in EVs samples (**Fig. 4a**). Additionally, EVVir samples were depleted in 3 ABC transporters (Atu1413, Atu3165, Atu2241), which could reflect a shift from the bacterial use of its EVs to competition. Finally, the virulence-induction condition had no significant effect on apolar and polar metabolites composition of EVs compared to the Glucose condition.

**Fig. 4:**
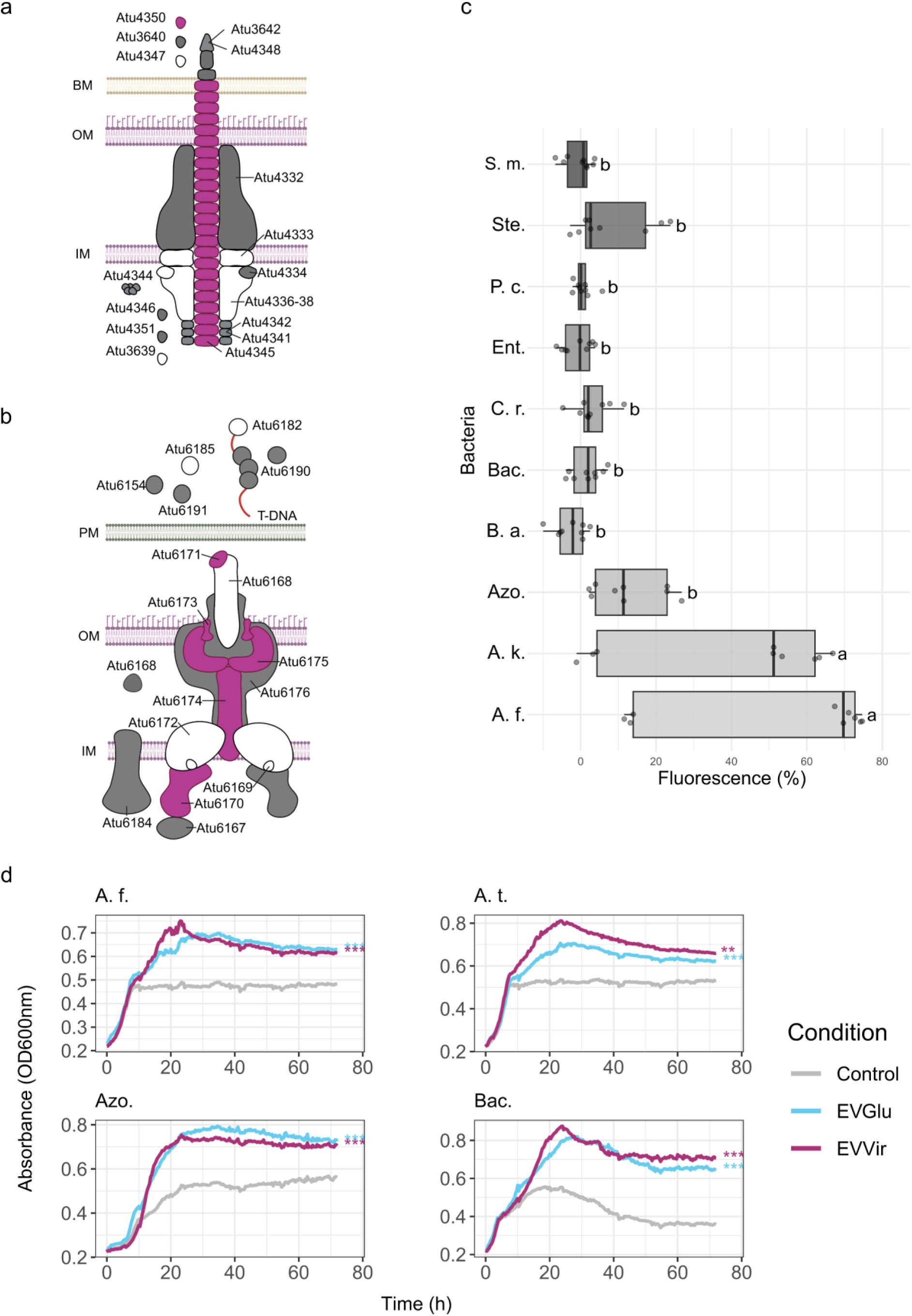
*A. fabrum* C58 EVs are involved in protein secretion and bacterial interactions. (**a**) Representation of the T6SS in *A. fabrum* C58. Proteins displayed are either structural proteins of the T6SS or involved in its assembly, effectors involved in bacterial competition and immunity proteins. (**b**) Representation of the T4SS in *A. fabrum* C58. Proteins displayed are either structural proteins of the T4SS or involved in its assembly, or effectors involved in tumor formation. Proteins enriched in EVs of *A. fabrum* C58 compared to the WCL in virulence condition are colored in purple. Proteins found in *A. fabrum* C58 EVs are colored in grey while absent proteins are colored in white. BM: bacterial membrane PM: plant membrane; OM: outer-membrane; IM: inner-membrane. (**c**) Association between *A. fabrum* C58 EVs and different bacterial strains cells. FM4-64 labeled EVs (1 mg.mL^-1^) from the EVGlu condition were incubated with bacterial cells for 1h at 28°C. Remaining EVs were quantified by calculating the remaining fluorescence in cell pellets (ex: 535 nm; ex: 615 nm) . Box plots show median (center line), interquartile range (box), and 1.5× interquartile range (whiskers) and technical replicates from 3 biological replicates shown as points. Statistical analysis was performed using the ANOVA with Tukey’s HSD post-hoc test, groups with different letters are statistically different (*p* < 0.05). (**d**) Most impacted bacterial growth by the presence of EVGlu or EVVir. Bacteria were co-inoculated with 5.10^8^ EVs particles from EVGlu and EVVir conditions in TSB 1/10 liquid medium for 72 h, *n* = 2 biological replicates. Statistical significance was determined by comparing the Area Under the Curve (AUC) for each group by ANOVA with a Tukey’s post-hoc test and the statistical differences between the Control and EVGlu or EVVir conditions are represented (**: *p* < 0.01; ***: *p* < 0.001). Bacterial strains A. f.: *Agrobacterium fabrum* C58, A. t.: *Agrobacterium tumefaciens* Kerr14, Azo.: *Azospirillum* sp. B510, B. a.: *Burkholderia ambifaria*; Bac.: *Bacillus sp.*; C. r.: *Chryseobacterium rhizosphaerae*; Ent.: *Enterobacter* sp.; P. c.: *Pseudomonas chlororaphis*; Ste.: *Stenotrophomonas* sp.; S. m.: *Serratia marescens*.

### Virulence induces the enrichment of toxins in *A. fabrum* C58 EVs compared to WCL

In virulence condition, EVs samples were enriched in 102 proteins and depleted in 60 proteins compared to WCL (**Fig. S5b**, **Table S2g**). Four proteins from the T6SS were found enriched in EV samples, from which two toxins Atu4345 (hcp) and Atu4350 (Tde1). Interestingly, the antitoxin Atu4351 abundance was not modulated in EVVir compared to WCL Vir (**Fig. 3a**). Atu1510, another putative toxin homolog to the toxin ParE4 of *Caulobacter vibrioides*, was also found enriched in EVVir samples compared to WCL Vir. As seen in Glu condition, EVVir were significantly enriched in periplasmic, outer-membrane and extracellular proteins while depleted in inner-membrane and cytoplasmic proteins (**Fig. S4**). Aside from proteins, EVVir samples were depleted in 26 and enriched in 20 apolar metabolites compared to the WCL (**Fig. S5d**, **Table S3g**). They were also depleted in 24 polar metabolites (**Fig. S5f**, **Table S4g**). Notably, EVs samples were depleted in amino acids and in fatty acids annotated as saturated and monounsaturated PE and PC species (**Fig. S5f**).

### *A. fabrum* C58 EVs carry many secretion system proteins

Despite not being significantly enriched in EVs compared to the WCL, many proteins belonging to T4SS and T6SS were found in EVVir samples. Regarding the T6SS, the structural proteins Atu4332 (TssM) and Atu4334 (TssK) that are both anchored to membranes, and Atu4341 (TssC41) and Atu4342 (TssB) that are components of the sheath were detected. The T6SS disassembly protein Atu4344 (TssH) was also found. In addition to structural proteins, many T6SS toxins were found like the three common T6SS toxins Atu4345 (TssD), Atu3642 (TssI2) and Atu4348 (TssI1). Notably, Atu4345 (TssD) was the 20th most abundant protein in EVs from the EVVir condition. Moreover, the excreted toxin Atu3640 (Tde2) was found without the antitoxin Atu3639 (Tdi2) (**Fig. 4a**). Concerning the T4SS, the structural proteins anchored the outer-membrane, Atu6173 (VirB7), Atu6174 (VirB8), Atu6175 (VirB9) and Atu6176 (VirB10) were found in EVVir samples. The cytoplasmic proteins Atu6170 (VirB4), Atu6177 (VirB11), Atu6184 (VirD4) and the extracellular Atu6171 (VirB5) and Atu6167 (VirB1) were also found. More importantly, the secreted effectors Atu6156 (VirF), Atu6190 (VirE2) and Atu6191 (VirE3) were also found in *A. fabrum* C58 EVs, with Atu6190 (VirE2) being the 101th most abundant protein in EVVir samples (**Fig. 4b**).

### *A. fabrum* EVs interact differently with environmental bacterial species

To test whether EVs from *A. fabrum* C58 could be relevant in bacterial interactions, we exposed *A. fabrum* C58 and 9 other bacterial strains frequently recovered in the rhizosphere to EVs from the EVGlu and EVVir conditions. The ability of EVs from the EVGlu or EVVir conditions to interact with bacteria was first evaluated in bacteria-EVs uptake assay using FM4-64 labeled EVs. Out of the 10 bacterial strains tested, *A. fabrum* C58 showed the best ability to uptake EVs from both the EVGlu and EVVir conditions. This uptake was not different from the uptakes of the closely related strain *Agrobacterium tumefaciens* Kerr14 (**Fig. 4c, Fig. S6**). Next, we tested the impact of EVs from both conditions on bacterial growth. Surprisingly, EVGlu and EVVir both significantly increased the growth of every strain but *Stenotrophomonas sp.* where only EVGlu increased its growth and *S. marescens* that was not impacted. Additionally, *A. fabrum* C58, *A. tumefaciens* Kerr14, *Azospirillum* sp. B510 and *Bacillus sp*. had their growth almost doubled compared to the Control condition (**Fig. 4d**, **Fig. S7**).

### *A. fabrum* C58 EVs content is internalized by the plant cells

Beyond interbacterial interaction, we then asked whether EVs cargo can be delivered inside plant cytosol. Given the abundance of VirE2 effector protein (Atu6190) inside EVVir, and in the light of approaches already done in the literature^38^, we decide to monitor VirE2 protein internalization by Bimolecular Fluorescent Complementation assay (BiFC). The split-GFP system consists of two non-fluorescent fragments, GFP1–10 and GFP11, which could bind each other spontaneously in order to restore the GFP fluorescence. To test protein delivery via bacterial EVs in A. thaliana cells, transgenic roots expressing a cytosolic sfGFP1–10 fragment were inoculated with FM4-64-labeled EVs produced in AB Virulence either by A. fabrum C58 WT or by the VirE2–sfGFP11 strain. Confocal imaging at 24 hpi showed no sfGFP signal above background in roots treated with WT EVs and in non-inoculated (NI) controls (**Fig. 5**). FM4-64 fluorescence was detected in EV-treated roots as rounded intracellular spots, predominantly within vascular cells, and was absent from NI roots. Traces of FM4-64 fluorescence signal were also detected in plant cell membranes. Strikingly, roots inoculated with EVs purified from the VirE2–sfGFP11 strain showed a clear increase in sfGFP fluorescence, indicating that the VirE2 protein was able to reach the plant cytosol (**Fig. 5**). Same observations were done in both wounded and unwounded plants. Together with our proteomic detection of Vir proteins in EVs from virulence conditions, these data indicate that EVs can transport virulence effectors to plant tissues and release them intracellularly.

**Fig. 5:**
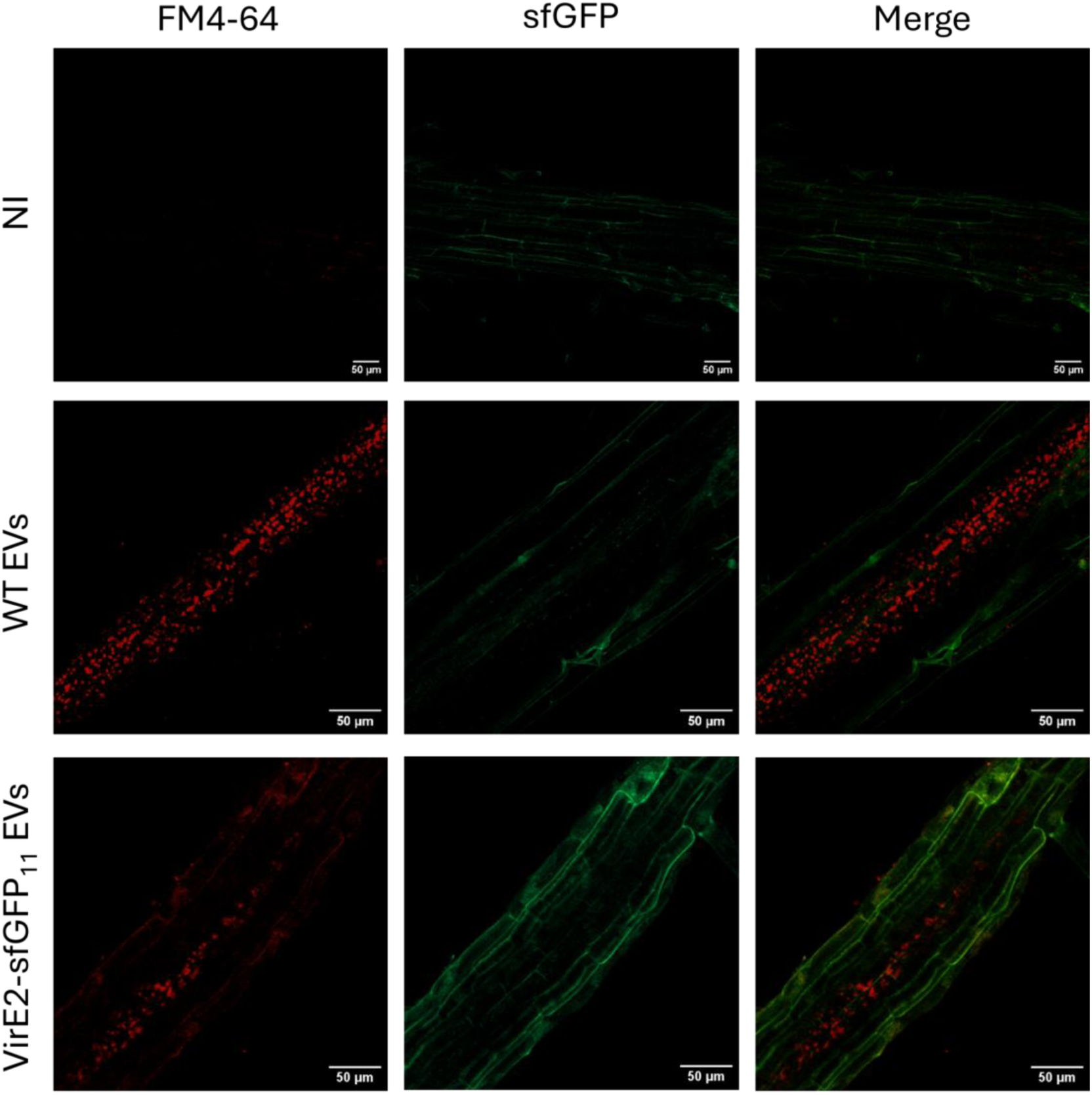
*A. fabrum* C58 EVs content is internalized by the plant. Representative images of wounded *A. thaliana* roots (24 hpi) expressing cytosolic sfGFP1–10 and inoculated or not with EVs purified from WT or VirE2-sfGFP11 expressing strains. Purified EVs were labeled with FM4-64 (1 mg.mL^-1^) prior to inoculation and images were acquired under identical settings (see Methods). Intracellular FM4-64 signal is visible in EV-treated roots, predominantly inside vascular cells, and specific sfGFP signal is observed in the EV VirE2–sfGFP11 condition only. *n* = 4 roots per condition per experiment; *n* = 3 biological replicates; ≥ 6 fields per root analyzed. NI = Non-inoculated.

### *A. fabrum* C58 EVs help in tumor formation

Given that EVs can mediate the transfer of virulence effectors to plant cells, we wondered if *A. fabrum* C58 EVs could contribute to tumor formation. Unsurprisingly, EVs alone are not able to induce cell proliferation as *S. lycopersicum* inoculated with EVs from whether the EVGlu or EVVir condition didn’t show any visible tumor as in the Control condition (**Fig. 6a**). This was confirmed by the absence of nopaline in their plant wound extracts. Nopaline is an opine produced by the plant only after T-DNA insertion in the host genome. However, co-inoculation with EVVir significantly enhanced the efficiency of tumor formation by *A. fabrum* C58 by increasing both their numbers and dry weight compared to tomato wounds inoculated with *A. fabrum* C58 alone (**Fig. 6a-c**). On the other end, the addition of EVGlu to *A. fabrum* C58 cells significantly increased the tumor colonization efficiency compared to *A. fabrum* C58 alone (**Fig. 6d**). Finally, EVs addition to *A. fabrum* C58 did not significantly impact the nopaline concentration determined within the tumor environment in either condition compared to *A. fabrum* C58 alone (**Fig. 6e**). Taken together with proteomics and imaging experiments, these results indicate that EVs increase the tumor formation efficiency and virulence of *A. fabrum* C58.

**Fig. 6:**
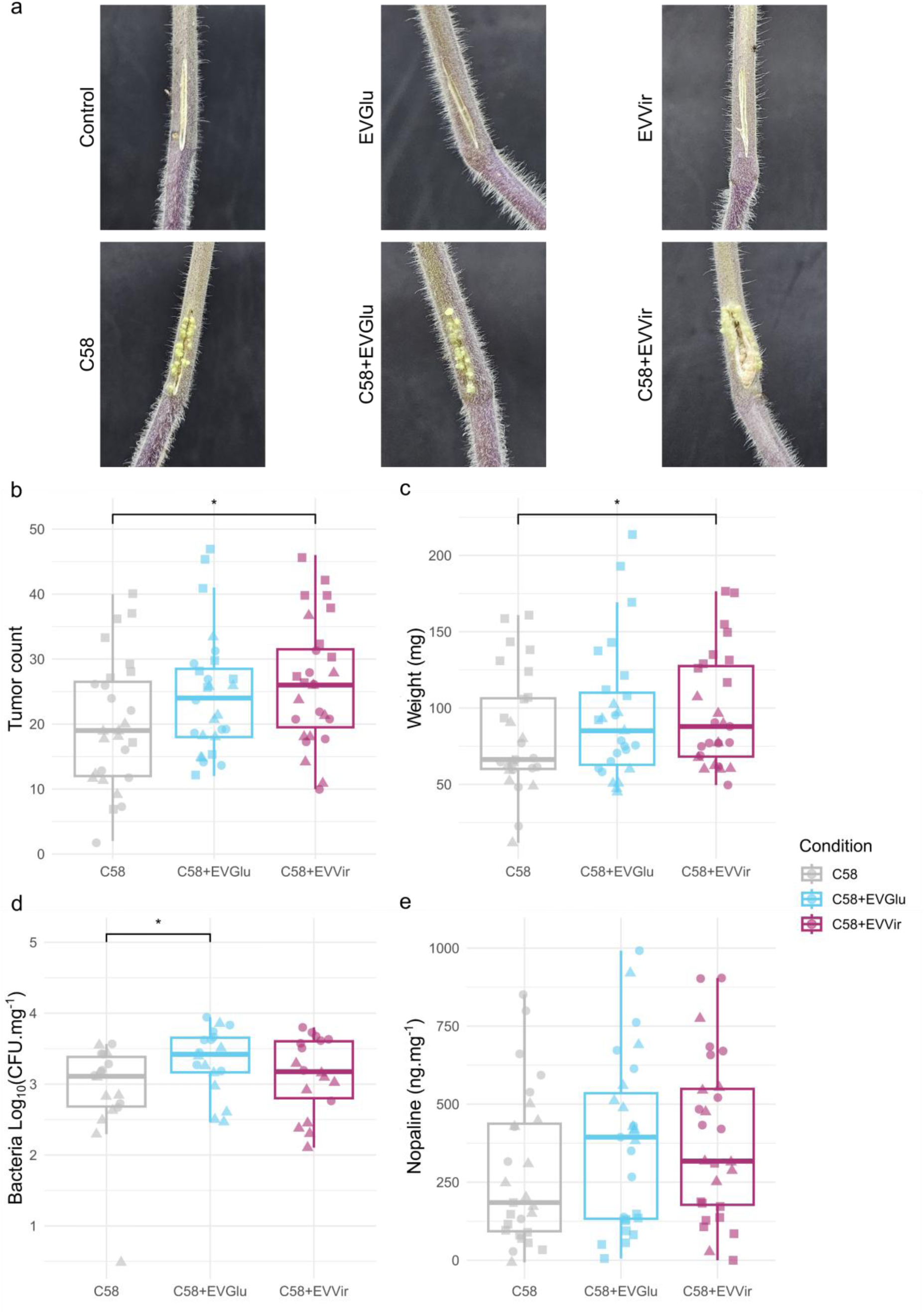
*A. fabrum* C58 EVs influence tumor formation in *S. lycopersicum*. (**a**) Pictures of wounded tomato stems 3 weeks after the inoculation of water as Control, EVGlu alone, EVVir alone, *A. fabrum* C58, *A. fabrum* C58 co-inoculated with EVGlu, *A. fabrum* C58 co-inoculated with EVVir. Different parameters were followed for tomato stems inoculated with *A. fabrum* C58 alone or supplemented with either EVGlu or EVVir : (**b**) tumor count, (**c**) dry weight of tumors, (**d**) number of *A. fabrum* C58 UFC in tumors and (**e**) nopaline concentration in tumors. The horizontal line within each box indicates the median, while the whiskers extend to the 1.5x interquartile range. Individual data points (*n* = 27) are overlaid to show biological variation, with shapes representing the three independent experimental blocks. Statistical significance was determined using a Linear Mixed-Effects Model with ’Condition’ as a fixed effect and ’Experimental block’ as a random effect, followed by Tukey’s HSD post-hoc test. Asterisks indicate significant differences between groups (*: *p* < 0.05).

### *A. fabrum* C58 EVs influences the plant root primary and specialized metabolites

As EVs from plant-associated bacteria have been shown to affect the plant’s response^16^, the influence of *A. fabrum* C58 EVs on the plant’s metabolome responses was analysed and compared with the impact of the living bacteria. Amongst the 24 tested amino acids in *S. lycopersicum* roots, the presence of EVs from *A. fabrum* C58 only decreased the concentration of histidine and threonine in the EVVir condition compared to the Control condition. However, *A. fabrum* C58 cells from the C58Glu condition increased the concentration of alanine, glycine and valine in tomato roots compared to the Control condition (**Fig. S8**). Untargeted metabolite analysis from the roots of *S. lycopersicum* led to the obtention of a data matrix containing 825 unique ions (**Table S5a**). PLS-DA analysis on the C58Glu, C58Vir and Control condition showed a separation between the three conditions, indicating that the bacteria impacted *S. lycopersicum* roots metabolites profiles differentially according to its cell state (**Fig. 7a**). It allowed to highlight the most discriminant ions between those conditions (385 ions, VIP score > 1). PLS-DA analysis performed on the EVGlu, EVVir and Control conditions indicated a clear separation between the EVVir and the Control condition. This separation was less clear when comparing EVGlu to the Control condition, mostly due to EVGlu variability. However, EVGlu and EVVir induced distinct metabolites profiles mostly due to the differential abundance of 423 ions (VIP score > 1) (**Fig. 7b**). Interestingly, both *A. fabrum* C58 cells and its purified EVs (in Glu or Vir condition) had no effect on *S. lycopersicum* aerial parts specialized metabolites (PLS-DA Q2Y scores < 1) or amino acids concentration (data not shown), indicating that only a local response was established in the plant 18h after inoculation.

**Fig. 7:**
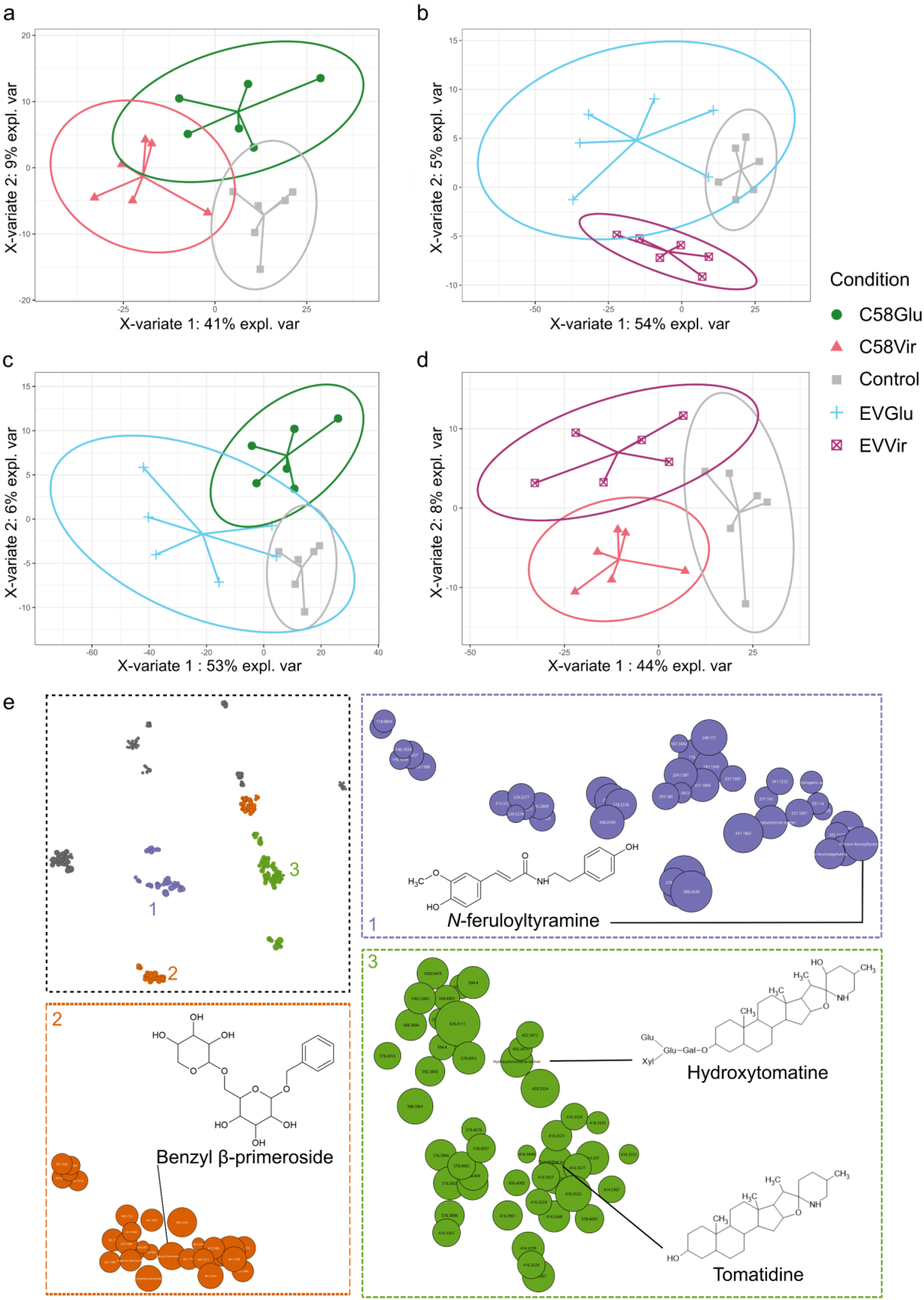
EVs from *A. fabrum* C58 influence the local abundance of plant defense metabolites 18 hpi. PLS-DA analyses on *S. lycopersicum* roots metabolite profiles comparing the Control condition (grey) to : (**a**) C58Glu (green) and C58Vir (red), (**b**) EVGlu (blue) and EVVir (purple), (**c**) C58Glu and EVGlu, (**d**) C58Glu and EVGlu. Each point represents the metabolite profile of one extract from pooled root samples of the same treatment (10 plants/pool). (**e**) t-SNE visualization of the molecular network obtained from*S. lycopersicum* root MS² data. Zoom-in clusters represent ions belonging to the same chemical family: (1, violet) hydroxycinnamic acid amides, (2, orange) O-glycoside compounds, (3, green) steroidal glycoalkaloids. Node size is proportional to the -Log10(*p*-value) of the explained difference between all conditions for each ion (*n* = 6, ANOVA with Tukey’s post-hoc test).

**Fig. 8:**
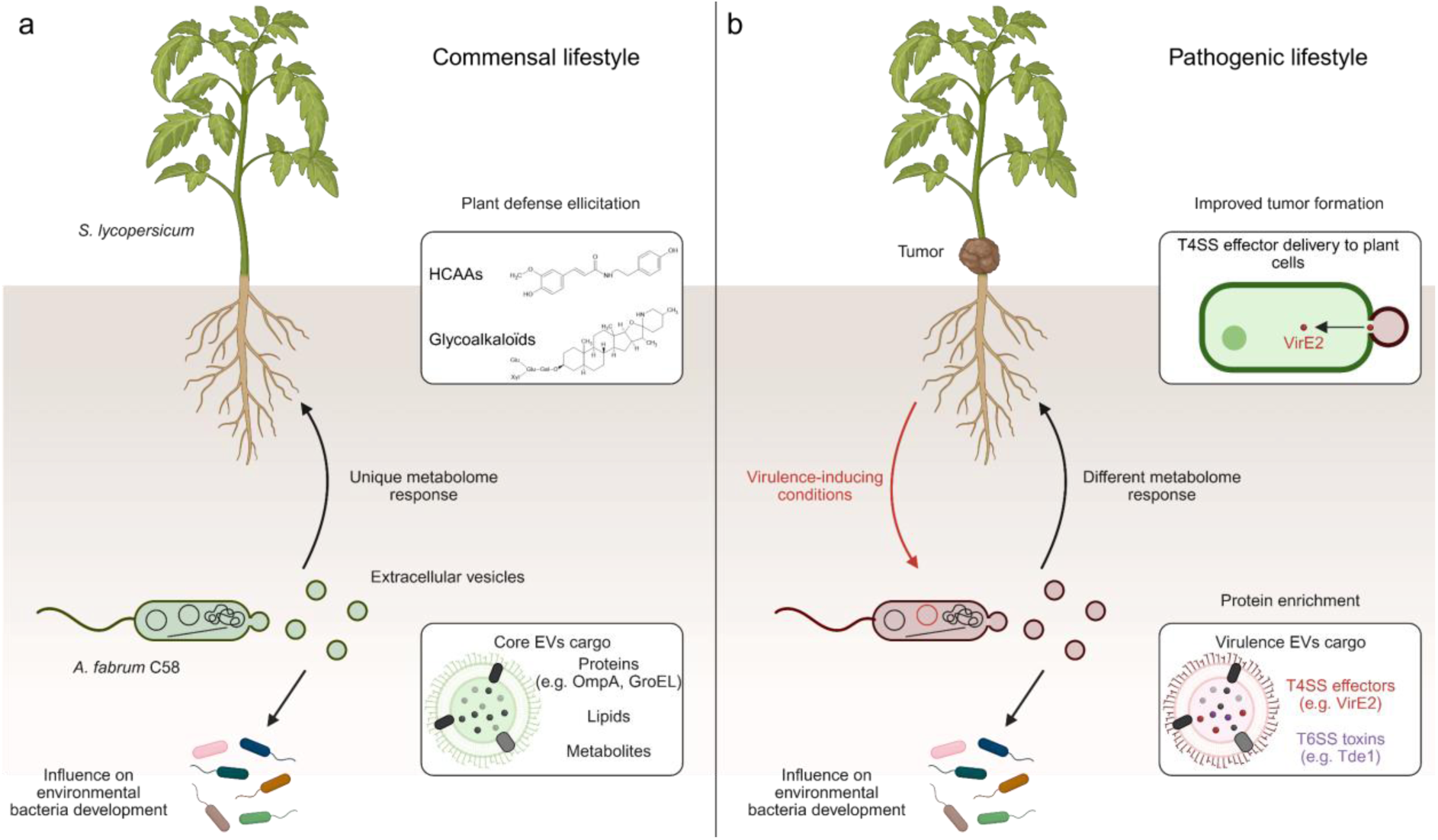
Multifaceted roles of EVs in *A. fabrum* C58 lifestyles. (**a**) In minimal growth conditions, *A. fabrum* C58 produces EVs with a cargo containing multiple proteins from which EVs markers, metabolites and lipids. These EVs can shape the bacterial community structure and induce a local metabolomic response in the plant host *Solanum licopersicum*. (**b**) In virulence-inducing conditions (*i.e*. glucose as carbon source, acetosyringone and acidic pH), *A. fabrum* C58 EVs are enriched in type IV and type VI secretion system proteins. EVs can also influence environmental bacteria development. On the plant side, EVs are able to fuse with plant cell membranes and deliver their cargo to the plant cell cytosol. Moreover, they enhance the tumor formation by *A. fabrum* C58 and trigger a different local metabolomic response than EVs from minimal condition.

### *A. fabrum* C58 EVs modulate the abundance of tomato defense metabolites

The UHPLC-UV/DAD-MS/MS QTOF data were explored to identify the discriminating compounds highlighted by statistical analyses. Study of the spectral data (UV-vis maxima; accurate mass; MS and MS/MS in positive and negative ionization mode) allowed the annotation and/or the identification of some compounds by comparison to databases, bibliographical data and analyses of standard compounds when available. Furthermore, to help identify series of key compounds involved in the response of *S. lycopersicum* roots to *A. fabrum* C58 and its EVs, MS² data were used to create a molecular network. The t-SNE visualization enables structurally related compounds to cluster together. Amongst the 825 detected unique ions, 648 ions were fragmented and 272 of them were assembled in 13 different clusters (**Fig. 7e**, **Table S5b**). Focusing on the clusters that showed the greatest differences in metabolite abundance between all conditions, three main clusters related to plant defense compounds were identified. In the first one, we annotated 5 hydroxycinnamic acid amides (HCAAs) derivating from ferulic acid: *N*-feruloylputrescine (C14H20N2O3, [M+H]^+^ at *m/z* 265.1547), *N*-feruloyl-lysine (C16H22N2O5, [M+H]^+^ at *m/z* 323.1603), *N*-feruloylagmatine (C15H19N3O3, [M+NH4]^+^ at *m/z* 307.1765), *N*-feruloyloctopamine (C18H19NO5, [M+H-H2O]^+^ at *m/z* 312.1230) and *N*-feruloyltyramine (C18H19NO4, [M+H]^+^ at *m/z* 314.1387). The second cluster corresponded to ions related to o-glycoside compounds from which benzyl ꞵ-primeveroside (C18H26O10, [M+NH4]^+^ at *m/z* 420.1877), violutoside (C19H26O12, [M+Na]^+^ at *m/z* 469.1319), isopentyl gentibioside (C17H32O11, [M+NH4]^+^ at *m/z* 430.2285), phenylethyl primeroside (C18H28O10, [M+NH4]^+^ at *m/z* 434.2023). Finally, the third cluster consisted of ions related to steroidal glycoalkaloids including tomatidine (C27H45NO2, [M+H]^+^ at *m/z* 416.3526), tomatine (C50H83NO21, [M+H]^+^ at *m/z* 1034.5539), hydroxytomatidine (C27H45NO3, [M+NH4]^+^ at *m/z* = 432.3474), hydroxytomatine (C50H83NO22, [M+H]^+^ at *m/z* 1050.5488), dehydrotomatine (C50H81NO21, [M+H]^+^ at *m/z* 1032.5379), ꞵ-1-tomatine (C45H75NO17, [M+H]^+^ at *m/z* 902.5118). Tomatine and tomatidine were confirmed by standard compound analyses. The MS² data of the identified or annotated compounds can be found in **Table S5c**.

### *A. fabrum* C58 EVs influence the plant defense metabolites differently than the bacteria

As EVs cargo differs from the whole cell content, EVs alone might impact differently the plant metabolome than the cell they originate from. Indeed, EVs from the Glucose condition impacted differently the root metabolite profile of *S. lycopersicum* than *A. fabrum* C58 whole cells and induced a more variable response (**Fig. 7c**). EVGlu had the strongest effect on defense metabolite abundances, inducing a significant increase in all annotated glycoalkaloids, O-glycosyl compounds and every HCAs and derivatives except the *N*-feruloyl-agmatine compared to the Control condition (**Fig. S9-11**). Moreover, it also induced a significant increase in 9 out of 10 annotated glycoalkaloids, 2 out of 4 annotated O-glycosyl compounds and 2 out of 9 annotated HCAs and derivatives compared to the C58Glu condition (**Fig. S9-11**). Interestingly, while PLS-DA analysis showed a separation between the EVVir, C58Vir and Control condition, none of the annotated defense metabolites except the *N*-feruloylagmatine showed a significant difference in accumulation between the EVVir and C58Vir conditions (**Fig. 7e**, **Fig. S9-11).** However, they were almost all significantly more abundant in these two conditions compared to the Control condition, indicating that specialized metabolites separating EVVir and C58Vir PLS-DA profiles might not be involved in the defense related metabolic pathways.

## Discussion

The environment plays a major role in phytobacteria ecology. Recent studies on EVs from plant-associated bacteria show that the production and cargo of bacterial EVs is modulated by plant cues^17^. Here, we showed that minimal or virulence-inducing conditions promote the production of EVs with different protein compositions in *A. fabrum* C58 and that these distinct EVs populations play different roles in *A. fabrum* C58 lifestyles, notably in tumor formation.

Regarding *A. fabrum* C58 EVs, our study showed that the protein cargo of EVs is composed of about 1600 unique proteins, a finding that is consistent with recent studies on bacterial EVs^39^. Most of the identified proteins in EVs were also found at high abundance in whole-cell lysates (WCL), indicating a possible “core” cargo composed of abundant proteins. However, specific virulence proteins were enriched in *A. fabrum* EVs when exposed to virulence-inducing conditions, suggesting an enrichment that reflects the physiological state of the bacteria in response to environmental changes. This idea is supported by *P. syringae* virulence effectors found both in EVs produced in virulence-inducing conditions and in EVs recovered *in planta* after infection^40^. Among the most abundant proteins in *A. fabrum* C58 EVs, we found several proteins related to T4SS and T6SS indicating their possible roles in interactions with other organisms, either with the host or microbial communities.

As bacterial secretion systems are anchored to membranes, it seems logical to think that their components and effectors are more likely to be loaded into vesicles. Firstly, the presence of secretion system components in membranes could induce a local deformation of the bacterial inner- and outer-membranes, thus inducing EVs formation at their location. Indeed, membrane proteins are known to induce changes in lipid membrane curvature^41^. Since the proteins associated with T4SS and T6SS are close to the secretion systems in *A. fabrum* C58, this could explain their high abundance if the bacterial secretion systems are a hotspot for EVs formation in cells^42^. Additionally, the presence of secretion systems at the bacterial membrane could induce a bias in the sorting and enrichment process of related proteins in the periplasm and EVs. Indeed, it has been shown recently that T6SS components allow the loading of cytoplasmic proteins lacking signal peptides into the EVs of the plant-associated bacteria *P. putida* KT2440, indicating the possible existence of multifunctional subcomplexes combining “classical” secretion systems and EVs^43^. Thus, the overabundance of T6SS proteins in *A. fabrum* C58 EVs purified from the virulence-inducing condition could explain the presence of cytoplasmic proteins in EVs. Moreover, this T6SS-mediated loading of cytoplasmic proteins could also happen with the T4SS as some of its components are also located in both inner-and outer-membranes.

In the plant environment, *A. fabrum* C58 interacts with multiple microbial species, especially bacteria. To overcome bacterial competition, *A. fabrum* C58 uses its T6SS to transfer toxins, such as DNAses, directly to the targeted cells cytoplasm^8^. Contrary to our expectations, EVs from the virulence-inducing conditions didn’t induce any killing but instead increased the growth of most of the challenged bacteria. Although it may seem counter intuitive, to date bacterial competition mediated by direct contact between EVs and bacterial cells was shown to be dependent of the presence of peptidoglycanases in the EVs cargo damaging the target cell integrity from the periplasmic space^44–46^. Here, the killing action of DNAses produced by *A. fabrum* C58 require them to be translocated to the cytoplasm of recipient cells and their transport within EVs might not allow them to reach this cellular compartment. Interestingly, *A. fabrum* C58 EVs were mostly uptaken by *A. fabrum* C58 itself or the closely related *A. tumefaciens* Kerr14. This specificity has been previously shown to be dependent on the presence of the outer-membrane protein Atu8019 in a liposome-bacteria context^35^. As this protein was found in both of our EVs population, it could also mediate the interaction between *Agrobacterium* sp. and their EVs. The fact that some bacteria species had their growth increased while they displayed low binding capacity to *A. fabrum* C58 EVs could highlight a system where EVs would represent a common good pool in the bacterial environment. These EVs could then be either absorbed by bacteria or would lyse in the extracellular environment, thus releasing nutrients. Moreover, bacteria such as *Bacillus* sp. could also actively lysate surrounding EVs to gain a privileged access to this nutrient source^47^.

*A. fabrum* C58 is mainly known for its ability to transfer genetic material, the T-DNA, and effectors to plant cells using T4SS machinery, thus inducing a hormonal imbalance leading to the creation of tumor due to cell proliferation^5,7^. We found in *A. fabrum* C58 EVs most of the T4SS proteins and virulence effectors, especially the virulence effector VirE2 at high abundance. Virulence effector presence in EVs has already been described in other phytopathogens and this study brings another example of EVs as an alternative to “classic” secretion systems^48^. Fusion of bacterial EVs to plant root cell membranes has recently been described between *Paraburkholderia phytofirmans* EVs and both *A. thaliana* or *S. lycopersicum*^49^. Here, we show that *A. fabrum* C58 EVs can also fuse to plant cell membranes. Membrane fusion of *Xanthomonas campestris* EVs to *A. thaliana* plant cells was shown to be dependent on EVs lipidic composition^50^. As EVs from the Glucose and Virulence condition didn’t show differences in lipid content, we can expect that both would interact the same way towards plant cell membranes but would not deliver the same cargo, thus inducing different plant responses. Interestingly, red-fluorescent spots were visualized inside root vessels (**Fig. 5**). In animal models, bacterial EVs have been shown to cross the intestinal epithelium either through or around epithelial cells to reach the blood stream^51^. However, the fate of bacterial vesicles in plants has not yet been studied, and the presence of these fluorescent structures raises questions: are they *A. fabrum* C58 EVs that have escaped fusion with plant membranes and reached the plant vessels? Or are they vesicles derived from plant membranes that have incorporated fluorescence after fusing with bacterial EVs? In any case, we show for the first time active virulence effector delivery from bacterial EVs to the plant cell cytosol, using VirE2 as a reporter. As VirE2 intracellular route has already been shown to be dependent on clathrin-mediated endocytosis followed by a travel along the endoplasmic reticulum / actin network toward the nucleus, it would be tempting to investigate if EV-delivered VirE2 follow the same pathways^52,53^. Altogether, these results support the importance of a better understanding of virulence factor, cargo delivery and traffic *via* EVs inside the host.

The involvement of *A. fabrum* C58 EVs in the bacterial pathogenicity was confirmed by their capacity to improve tumor formation and colonization efficiency. However, they are not sufficient to induce tumor formation on their own and we propose that they act as helpers by delivering virulence effectors while also impacting plant physiology in other ways. Indeed, previous studies show that EVs from phytobacteria are able to elicit immune responses in plants, mostly through the induction of genes related to hormonal signalisation and defense pathways^16^. *A. fabrum* C58 is mainly studied in the context of crown-gall disease, but the bacteria also evolves in the soil and rhizosphere of many plants^4^. Using untargeted LC-MS² analysis, we show that EVs from *A. fabrum* C58 can modulate the local defense response in *S. lycopersicum* roots. Interestingly, EVs from *A. fabrum* C58 induced the accumulation of hydroxycinnamic acid amides and glycoalkaloids, two defense-related metabolites families^54,55^. These classes of molecules were previously shown to also be accumulated in presence of EVs from the phytobeneficial bacteria *Azospirillum* sp. B510^28^. These results could indicate a common plant response to EVs derived from rhizospheric bacteria. Moreover, we show for the first time that EVs induce a specific response in plants compared to the bacteria they originate from. Despite the use of a multiomic approach, these differences could not be only attributed to compounds found differentially accumulated in *A. fabrum* C58 EVs compared to the cells. However, it is now well established that the plant response to bacterial EVs is complex and involves multiple molecular actors^16,56^. Additionally, our study did not investigate the nucleic acid content of *A. fabrum* C58 EVs. The nucleic acids, especially small interfering RNA (sRNAs), could be involved in the modulation of *S. lycopersicum* physiology. Indeed, EVs from the phytopathogenic *Xylella fastidiosa* were shown to transport sRNAs that target and lower the expression of defense-related genes in *A. thaliana*^57^.

In conclusion, our work reveals that the plant pathogen *Agrobacterium fabrum* C58 utilizes extracellular vesicles as a dynamic and multifaceted platform for inter-organismal communication. We have demonstrated that EVs are not merely passive carriers but are actively loaded with virulence effectors and toxins in response to environmental cues, enabling them to enhance pathogenicity and influence the growth of environmental bacteria. This study establishes that EVs play a key role in bacterial virulence and adaptation, providing new insights into how bacteria interact with their host and surrounding microorganisms.

## Acknowlegments

A part of this work was financially supported by the ANR program (project PhyEVE), the French national program EC2CO (project InteractOMVs) and by the BioEEnViS Research Federation (project VEX). TZP received a doctoral grant from the French Ministère de l’Education Nationale, de l’Enseignement Supérieur et de la Recherche. We acknowledge the DTAMB platform of BioEEnViS Research Federation for providing access to the ultracentrifuge; the CTµ (Centre Technologique des Microstructures, Université Lyon 1) for the electron microscopy and confocal observations; the “Serre et chambre climatiques” platform of BioEEnViS Research Federation for tomato culture; the Centre d’Etudes des Substances Naturelles (UMR 5557 Ecologie Microbienne) for their help and recommendations on the metabolomic experiment, analysis and fruitful discussions (M. Rey, G. Meiffren, P-E Mercier, P. Vergne and T. Herrera Durigneux). We acknowledge A. Page from Protein Science Facility of SFR Biosciences (UAR3444/CNRS, US8/Inserm, ENS de Lyon, Université Lyon 1) for mass spectrometry analysis. We acknowledge I. Tguafaiti (Université Lyon 1, CNRS, INSA, MAP, UMR 5240, BAYER) for kindly providing access to the NTA. We also acknowledge M. Pallier for its skillful technical assistance in plant metabolites extractions. We would like to thank B. Schaack for providing liposomes stocks and Karl-Erich Jaege for sharing sfGFP1-10 coding sequence. Graphical figures were created using <u>Biorender.com</u>.

## Author contribution

T. Z-P, C. L and L. V. designed the study. T. Z-P, D. S. and F. N. isolated vesicles. T. Z-P and F. N. performed the NTA analysis and T. Z-P performed the TEM image acquisition. T. Z-P performed the EVs and bacterial content extraction and multi-omics analysis with the help of V. G., I. K. and G. C. D. S. and J. D. performed the bacterial challenge and binding experiments. F. N. conducted the confocal imagery experiment with the help of D. S. and F. X-G. Tumor experiments were done by T. Z-P, D. S., F.N., C. L., L.V., V. G. and I.K. Plant metabolites experiment was conducted by T. Z-P and V. G., I.K. and G.C. helped with the analysis. T. Z-P. wrote the first version of the manuscript and prepared the figures. F.N., F. X-G., G. C., I. K., C. L. and L. V. contributed to the final manuscript writing. All the authors approved the final version of the manuscript.

## Supplementary tables

**Table S1:** Bacterial strains and PCR primers

**Table S2:** Proteomic analysis of *A. fabrum* C58 EVs and WCL

**Table S3:** Lipidomic analysis of *A. fabrum* C58 EVs and WCL

**Table S4:** Metabolomic analysis of *A. fabrum* C58 EVs and WCL

**Table S5:** Impact of *A. fabrum* C58 and its EVs on *S. lycopersicum* root metabolome

## Supplementary figures

**Fig. S1:** Virulence induction of *A. fabrum* C58 in AB Glucose and AB Virulence.

**Fig. S2:** *A. fabrum* C58 produces OMVs by blebbing from the outer-membrane.

**Fig. S3:** *A. fabrum* C58 EVs purification by GDUC.

**Fig. S4:** *A. fabrum* C58 EVs from the virulence condition are enriched in OM proteins.

**Fig. S5:** Molecular enrichment in EVGlu and EVVir compared to *A. fabrum* C58 WCL of the bacteria they originate from.

**Fig. S6:** *A. fabrum* C58 EVVir interacts mostly with closely related species.

**Fig. S7:** *A. fabrum* C58 EVs affect the growth of other environmental bacteria.

**Fig. S8:** *S. lycopersicum* roots amino acids under the influence of *A. fabrum* C58 and its EVs.

**Fig. S9:** Annotated glycoalkaloids of *S. lycopersicum* roots impacted by *A. fabrum* C58 and its EVs.

**Fig. S10:** Annotated hydroxycinnamic acids and derivatives of *S. lycopersicum* roots impacted by *A. fabrum* C58 and its EVs.

**Fig. S11:** Annotated O-glycoside compounds of *S. lycopersicum* roots impacted by *A. fabrum* C58 and its EVs.

## Supplementary text

**1.** AB medium and virulence-inducing AB medium composition
**2.** MPLEx protocol
**3.** Protein analyses
**4.** *A. fabrum* C58 polar and apolar metabolites analyses
**5.** Extraction of nopaline from tomato tumors
**6.** Extraction of *S. lycopersicum* metabolites
**7.** Targeted analysis of free amino acid content of *S. lycopersicum*
**8.** Analysis of specialized metabolites content of *S. lycopersicum*
**9.** Metabolite annotation and molecular networking
**10.** Statistical analysis

