## Supplementary figures for "Multifaceted roles of extracellular vesicles in *Agrobacterium fabrum* C58 lifestyles"

#### Authors and affiliations

Timothée ZANNIS-PEYROT (1), Fanny NAZARET (2), Deniz SARIGOL (3), Jeanne DORE (4), François-Xavier GILLET (5), Vincent GAILLARD (6), Gilles COMTE (7), Isabelle KERZAON (8), Céline LAVIRE (9)(*), Ludovic VIAL (10)(*)(**)

* Both authors participated equally to this work

** Corresponding author

### Supplementary Figures

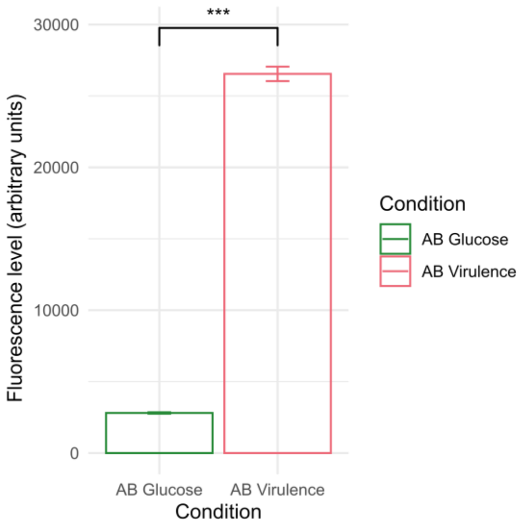


**Fig. S1: Virulence induction of *A. fabrum* C58 in AB Glucose and AB Virulence.**

*virB* expression (transcriptional fusion between *virB* promoter and eGFP) by *A. fabrum* C58 in AB Glucose or AB Virulence growth medium after 31 h. The fluorescence level (in arbitrary units) was corrected by dividing the fluorescence value by the $OD_{600nm}$. Results show the mean ± SD of fluorescence (ex: 485 nm; em: 530 nm) normalized by $OD_{600nm}$ ($n = 3$ biological replicates, two-sided Student's *t*-test, *** : $p < 0.001$).

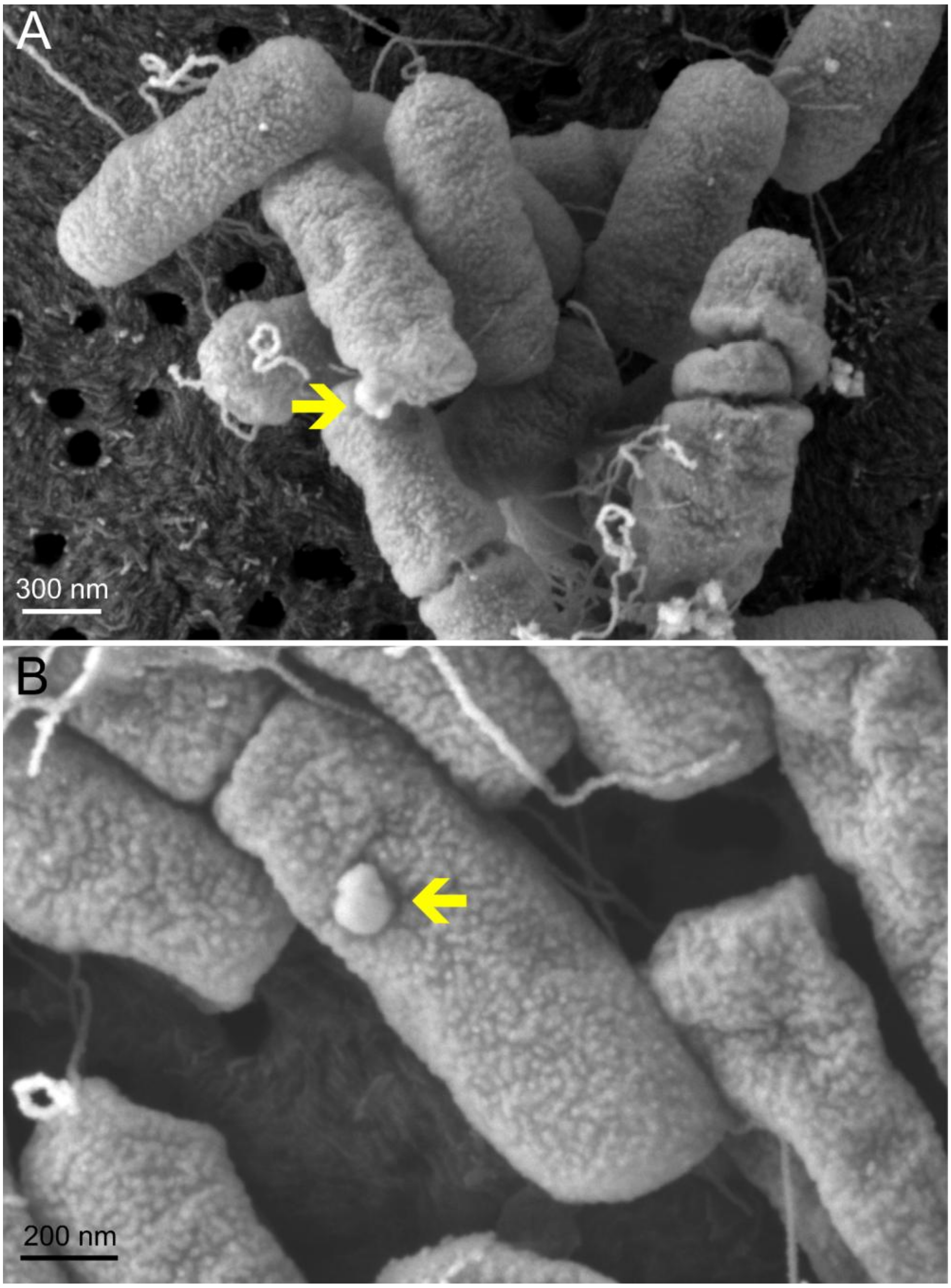


**Fig S2:** ***A. fabrum*** **C58 produces OMVs by blebbing from the outer-membrane.**

Scanning electron micrographs of *A. fabrum* C58 cells and EVs blebbing from the outer-membrane. Yellow arrows indicate the presence of EVs.

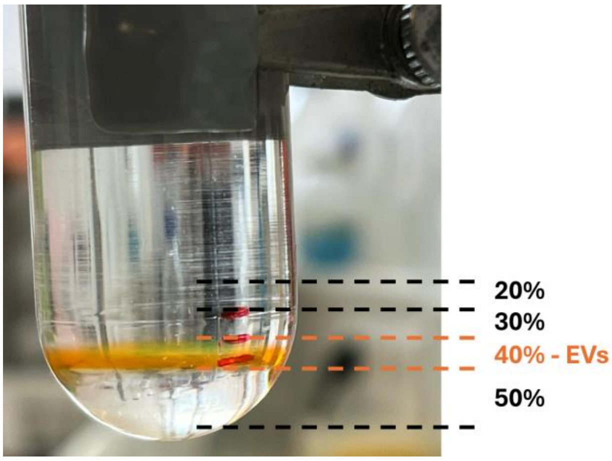


**Fig. S3: *A. fabrum* C58 EVs purification by GDUC.**

Picture of Optiprep gradient-density after migration of EVs labelled with DiO lipophilic dye (1 mg.mL$^{-1}$). EVs migrate to the 40% layer.

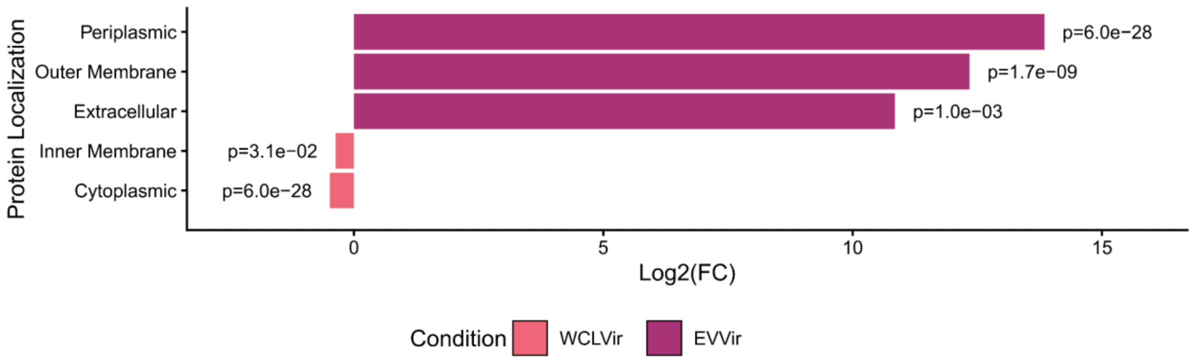


**Fig. S4: *A. fabrum* C58 EVs from the virulence condition are enriched in OM proteins.**

Comparison of the protein predicted localization between EVVir and WCLVir. Protein localization was predicted using DeepLocPro 1.0. Enrichment was calculated using Fisher's test between the two compared conditions. Protein localizations were considered enriched or depleted when $p < 0.05$.

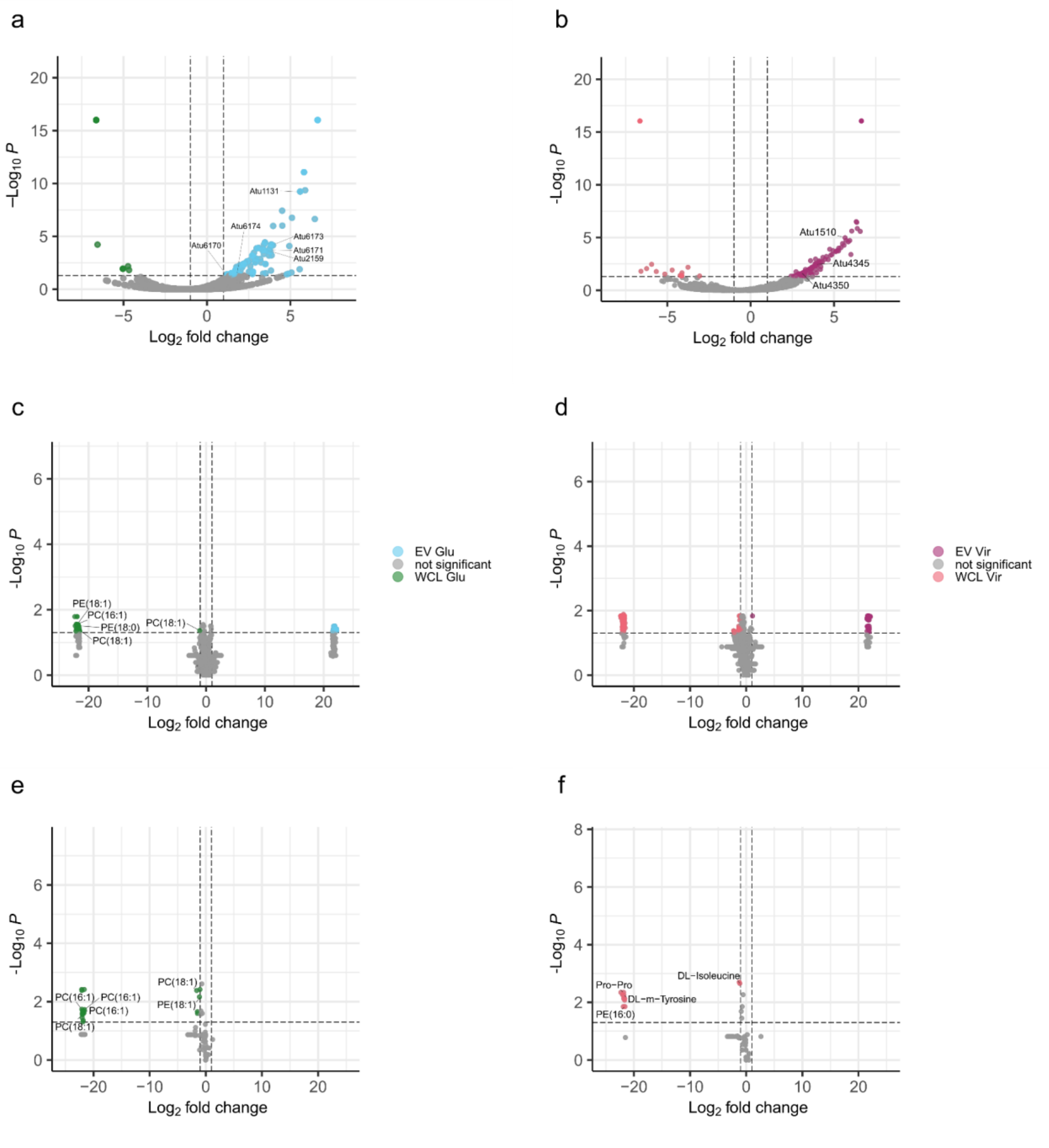


**Fig. S5: Molecular enrichment in EVGlu and EVVir compared to *A. fabrum* C58 WCL of the bacteria they originate from.**

Protein, apolar and polar metabolites content was separated following MPLEx protocol on 3 biological replicates. Volcano plot displaying differential abundance of (**a**) protein, (**c**) apolar metabolites, (**e**) polar metabolites between EVGlu (right, cyan) and WCLGlu (left, green). Volcano plot displaying differential abundance of (**b**) protein, (**d**) apolar metabolites, (**f**) polar metabolites between EVVir (right, purple) and WCLVir (left, red). Differential abundance was determined by a Student's *t*-test with BH correction ($p < 0.05$, $Log_2$-fold change > |0.5|).

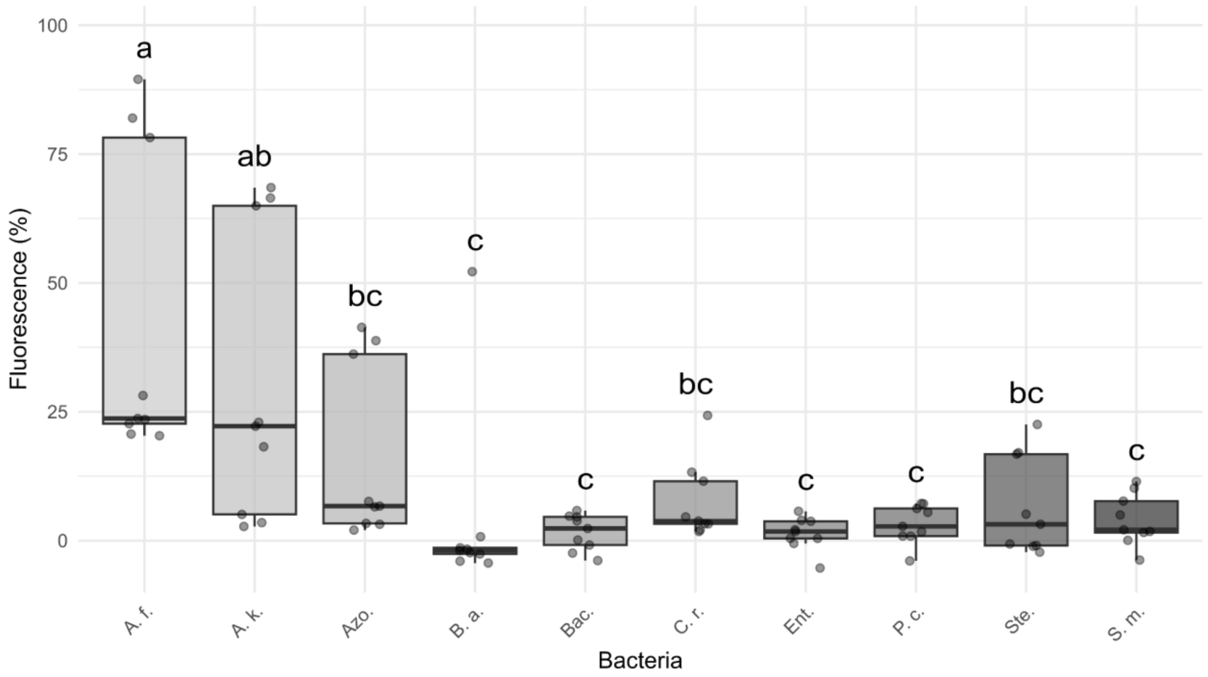


**Fig. S6: *A. fabrum* C58 EVVir interacts mostly with closely related species.**

Association between *A. fabrum* C58 EVs and different bacterial strains cells. FM4-64 labeled EVs (1 mg.mL$^{-1}$) from the EVVir condition were incubated with bacterial cells for 1 h at 28°C. Remaining EVs were quantified, calculating the remaining fluorescence (ex: 535 nm; ex: 615 nm). Box plots show median (center line), interquartile range (box), and 1.5× interquartile range (whiskers) and technical replicates from $n$ = 3 biological replicates shown as points. Statistical analysis was performed using the ANOVA with Tukey's HSD post-hoc test, groups with different letters are statistically different ($p < 0.05$).

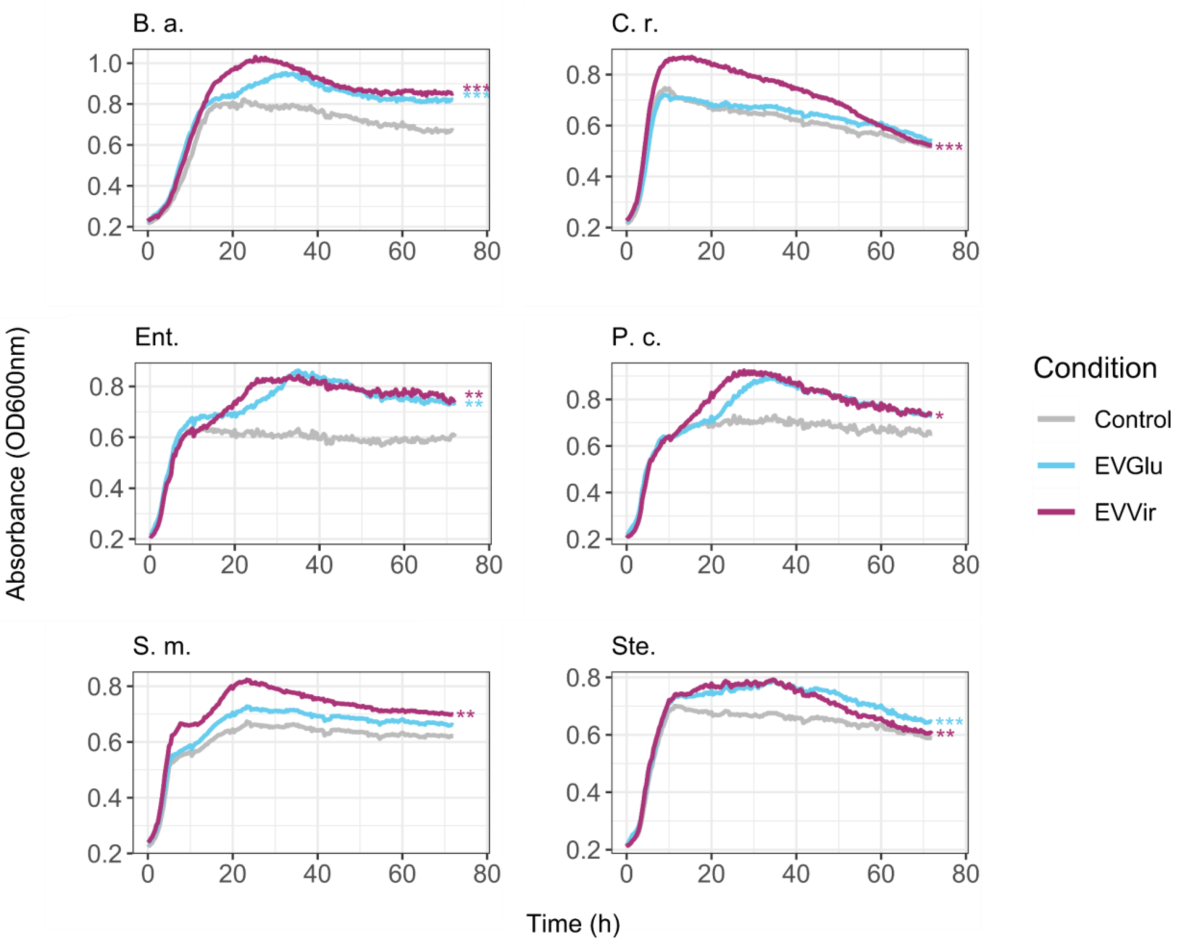


**Fig. S7: *A. fabrum* C58 EVs affect the growth of other environmental bacteria.**

Impact on bacterial growth of the presence of EVGlu or EVVir. Bacteria were co-inoculated with $5.10^8$ EVs particles from EVGlu and EVVir conditions in TSB 1/10 liquid medium for 72 h, *n* = 2 biological replicates. Statistical significance was determined by comparing the Area Under the Curve (AUC) for each group with by ANOVA with a Tukey's post-hoc test and the statistical differences between the Control and EVGlu or EVVir conditions are represented (*: $p < 0.05$; **: $p < 0.01$; ***: $p < 0.001$). All strains but *S. m.* and *Ste.* were significantly impacted by both EVGlu and EVVir presence. B. a.: *Burkholderia ambifaria*; C. r.: *Chryseobacterium rhizosphaerae*; Ent.: *Enterobacter* sp.; P. c.: *Pseudomonas chlororaphis*; Ste.: *Stenotrophomonas* sp.; S. m.: *Serratia marescens*.

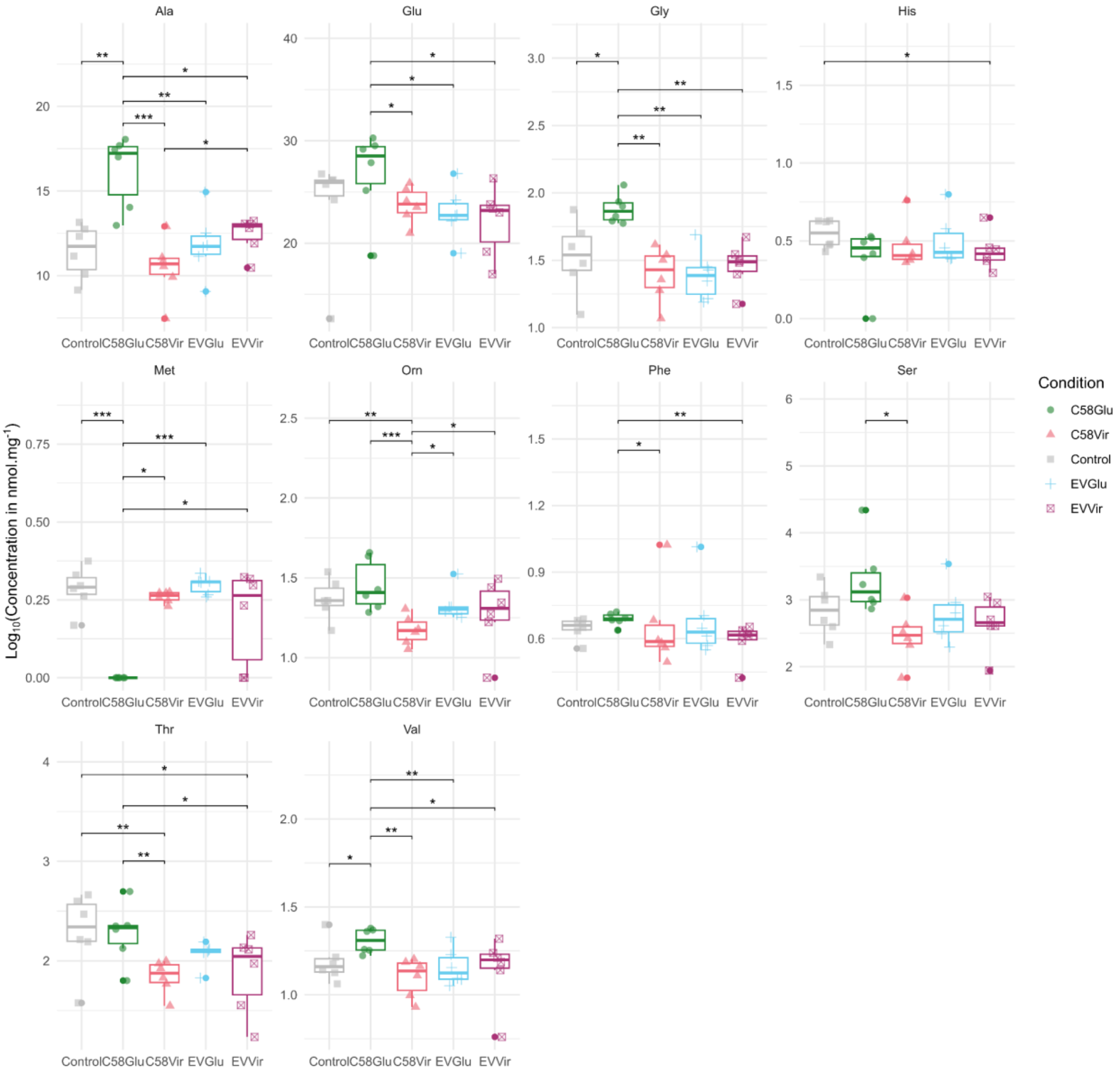


**Fig. S8: *S. lycopersicum* roots amino acids under the influence of *A. fabrum* C58 and its EVs.**

Amino acids concentration was determined using HPLC-UV/DAD-FLD after inoculation of EVs (EVGlu and EVVir), *A. fabrum* C58 (C58Glu and C58Vir) cells or water (Control) conditions on the roots of *S. lycopersicum*. Amino acids were quantified 18 hpi. Ala: alanine; Glu: glutamine; Gly: glycine; His: histidine; Met: methionine; Orn: ornithine; Phe: phenylalanine; Ser: serine; Thr: threonine; Val: valine. Box plots show median (center line), interquartile range (box), and 1.5× interquartile range (whiskers) of each amino acid concentration $Log_{10}(nmol.mg^{-1})$ ($n = 6$ biological replicates, 10 plants/pool for each replicate, ANOVA with Tukey's post-hoc test, *: $p < 0.05$, **: $p < 0.01$, ***: $p < 0.001$).

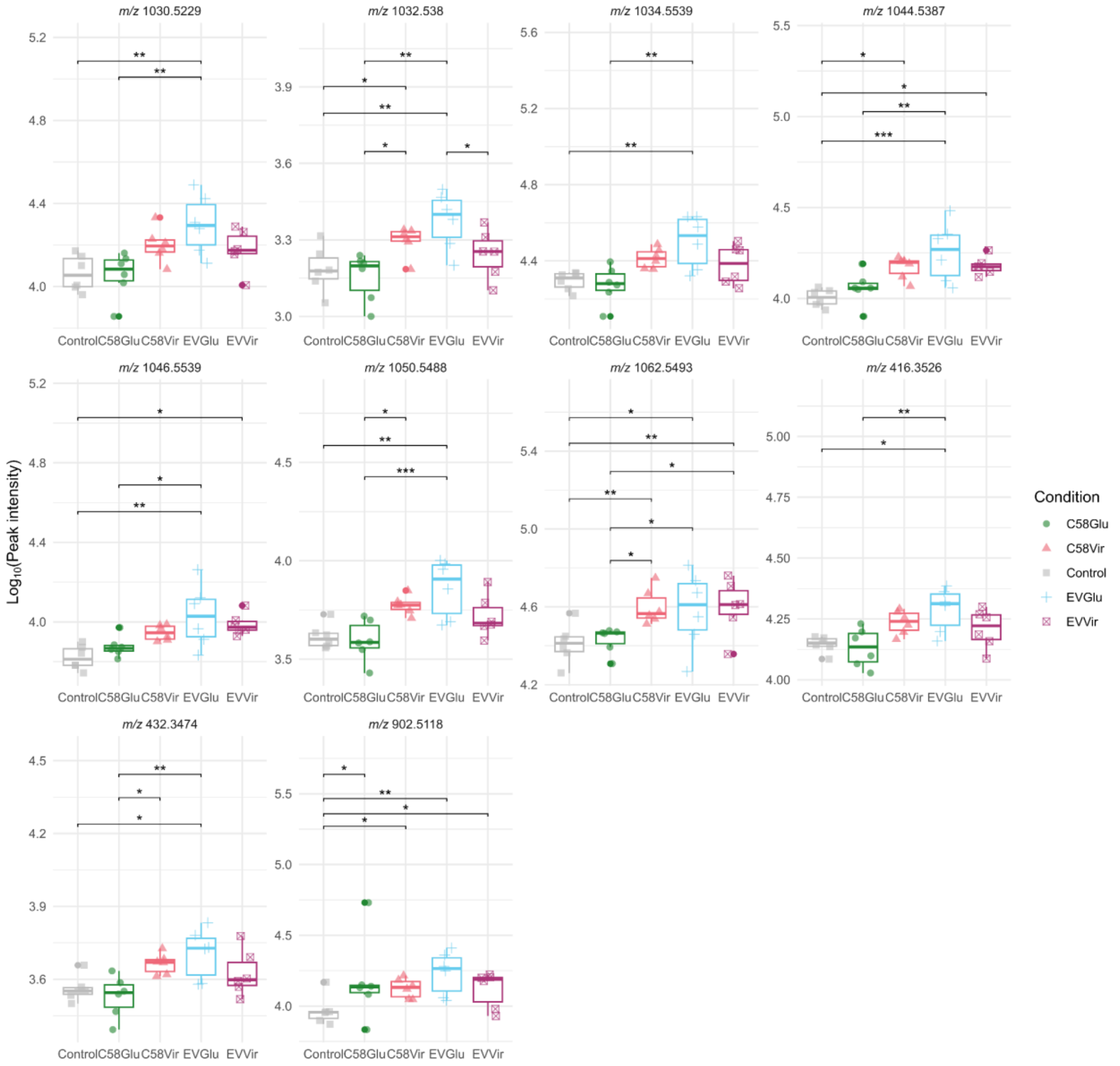


**Fig. S9: Annotated glycoalkaloids of *S. lycopersicum* roots impacted by *A. fabrum* C58 and its EVs.**

Impact of EVs (EVGlu and EVVir), *A. fabrum* C58 (C58Glu and C58Vir) cells or Control condition on the glycoalkaloid content of *S. lycopersicum* roots 18 hpi. *m/z* = 416.3526: tomatidine; *m/z* = 432.3474: hydroxytomatidine isomer; *m/z* = 902.5118: β-1-tomatine isomer; *m/z* = 1030.5229: unidentified glycoalkaloid; *m/z* = 1032.5379: dehydrotomatine isomer; *m/z* = 1034.5539: tomatine; *m/z* = 1044.5387: unidentified glycoalkaloid; *m/z* = 1046.5539: unidentified glycoalkaloid; *m/z* = 1050.5488: hydroxytomatine isomer; *m/z* = 1062.5493: unidentified glycoalkaloid. Box plots show median (center line), interquartile range (box), and 1.5× interquartile range (whiskers) interquartile range (box), 1.5× interquartile range (whiskers) of each ion's $\text{Log}_{10}$(peak intensity) ($n = 6$ biological replicates, 10 plants/pool, ANOVA with Tukey's post-hoc test, *: $p < 0.05$, **: $p < 0.01$, ***: $p < 0.001$).

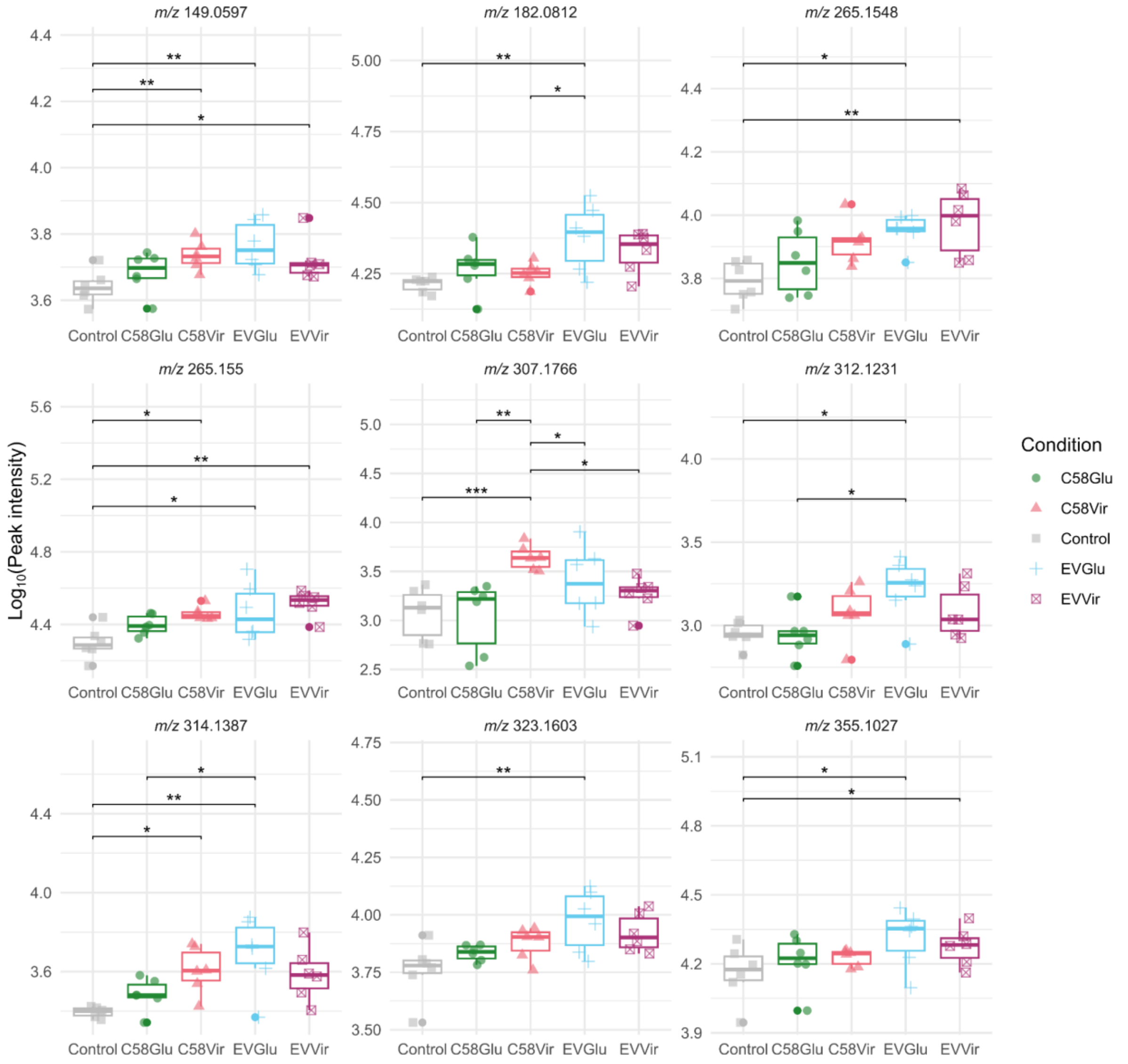


**Fig. S10: Annotated hydroxycinnamic acids and derivatives of *S. lycopersicum* roots impacted by *A. fabrum* C58 and its EVs.**

Impact of EVs (EVGlu and EVVir), *A. fabrum* C58 (C58Glu and C58Vir) cells or Control condition on the hydroxycinnamic acids and derivatives content of *S. lycopersicum* roots 18 hpi. *m/z* = 149.0597: 2-hydroxycinnamaldehyde; *m/z* = 182.0812 : coumaric acid; *m/z* = 265.1548: *N*-feruloylputrescine isomer; *m/z* = 265.1550 : *N*-feruloylputrescine isomer; *m/z* = 307.1766: *N*-feruloylagmatine; *m/z* = 312.1231: *N*-feruloyloctopamine; *m/z* = 314.1387 : *N*-feruloyltyramine; *m/z* = 323.1603: *N*-feruloyl-lysine; *m/z* = 355.1027: chlorogenic acid. Box plots show median (center line), interquartile range (box), and 1.5× interquartile range (whiskers) interquartile range (box), 1.5× interquartile range (whiskers) of each ion's $\text{Log}_{10}$(peak intensity) (*n* = 6 biological replicates, 10 plants/pool, ANOVA with Tukey's post-hoc test, *: $p < 0.05$, **: $p < 0.01$, ***: $p < 0.001$).

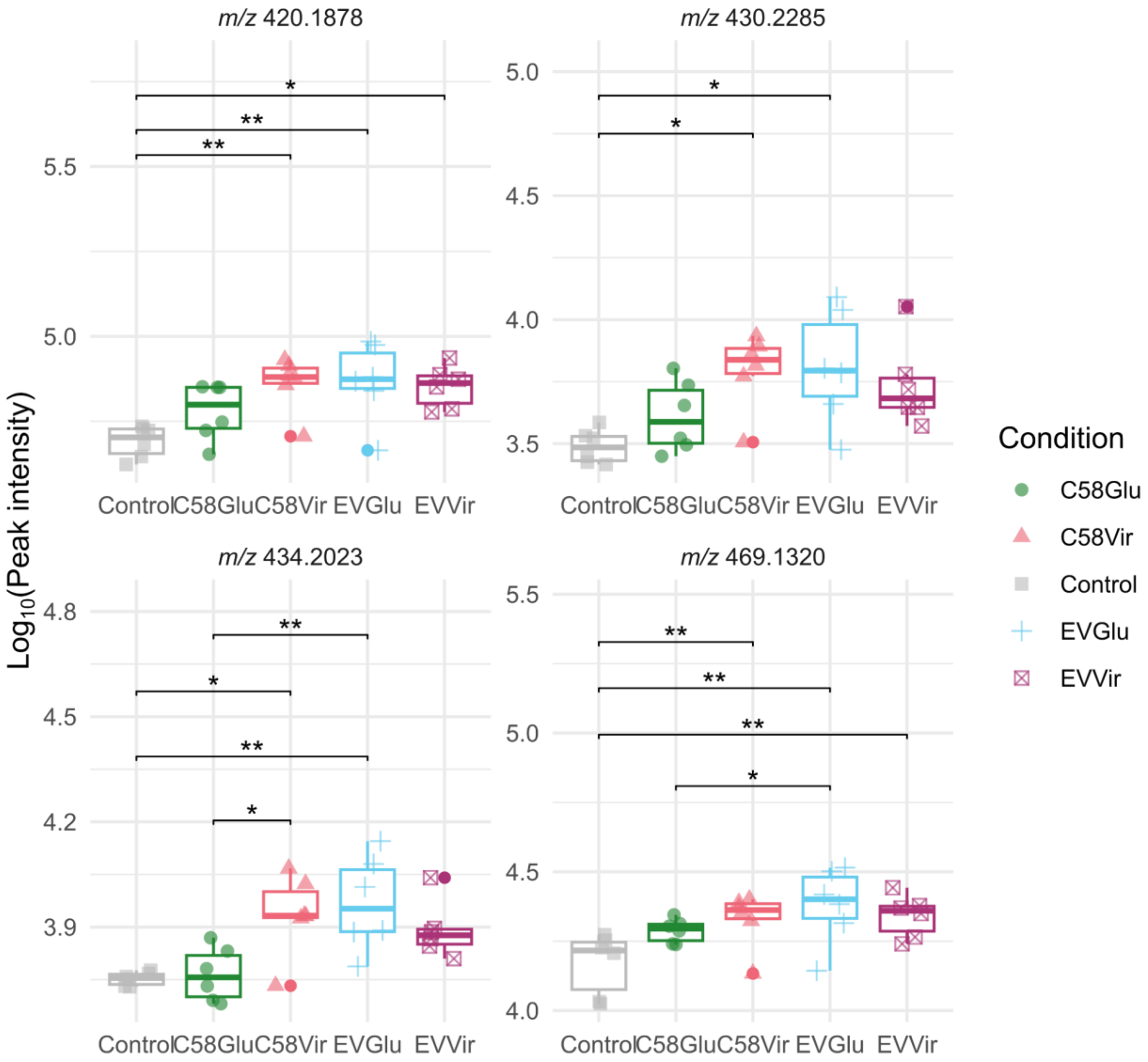


**Fig. S11: Annotated O-glycoside compounds of *S. lycopersicum* roots impacted by *A. fabrum* C58 and its EVs.**

Impact of EVs (EVGlu and EVVir), *A. fabrum* C58 (C58Glu and C58Vir) cells or Control condition on the O-glycoside compounds content of *S. lycopersicum* roots 18 hpi. *m/z* = 420.1878: benzyl β-primeveroside; *m/z* = 430.2285: violutoside isomer; *m/z* = 434.2023: isopentyl gentibioside; *m/z* = 469.1320: phenylethyl primeveroside. Box plots show median (center line), interquartile range (box), and 1.5× interquartile range (whiskers) interquartile range (box), 1.5× interquartile range (whiskers) of each ion's $Log_{10}$(peak intensity) ($n = 6$ biological replicates, 10 plants/pool, ANOVA with Tukey's post-hoc test, *: $p < 0.05$, **: $p < 0.01$, ***: $p < 0.001$).
