## Supplementary text for "Multifaceted roles of extracellular vesicles in *Agrobacterium fabrum* C58 lifestyles"


#### Authors and affiliations

Timothée ZANNIS-PEYROT (1), Fanny NAZARET (2), Deniz SARIGOL (3), Jeanne DORE (4), François-Xavier GILLET (5), Vincent GAILLARD (6), Gilles COMTE (7), Isabelle KERZAON (8), Céline LAVIRE (9)(*), Ludovic VIAL (10)(*)(**)

* Both authors participated equally to this work

** Corresponding author

### Supplementary Text

#### 1. AB medium and virulence-inducing AB medium composition

AB minimal medium was assembled by adding 50 mL of AB salt stock solution (20X; $NH_4Cl$ 20 g, $MgSO_4$ .$7H_2O$ 6 g, KCl 3 g, $CaCl_2$ 200 mg, $FeSO_4.7H_2O$ 50 mg, $dH_2O$ 1 L), 50 mL of AB buffer stock solution (20X; $K_2HPO_4$ 60 g, $NaH_2PO_4$ 23 g, $dH_2O$ 1 L) and 100 mL of glucose 100 mM (final concentration 10 mM) to 800 mL of sterile $dH_2O$. Virulence-inducing AB medium was assembled by adding 50 mL of AB salt stock solution (20X), 1 mL of AB buffer stock solution (20X), 5 mL of phosphate 0.1 M pH 7.0 buffer solution ($NaH_2PO_4.7H_2O$ 20.2 g, $NaH_2PO4.H_2O$ 3.4 g, $dH_2O$ 1 L), 100 mL of 2-(N-Morpholino)ethanesulfonic acid hydrate, 4-Morpholineethanesulfonic acid (MES) 0.5 M pH 5.5 buffer solution (97.6 g, $dH_2O$ 1 L) and 100 mL of glucose 100 mM (final concentration 10 mM) to 800 mL of sterile $dH_2O$ supplemented with acetosyringone (final concentration 15 µM).

#### 2. MPLEx protocol

Metabolites, proteins and lipids from 3 biological replicates of *A. fabrum* C58 cells or their EVs purified from cell cultures in AB Glucose or AB Virulence medium were extracted according to **Nakayasu et al., 2016**[1] with following adjustments. EVs samples or *A. fabrum* C58 cells were resuspended in 1 mL of MS-grade $H_2O$. Five milliliters of cold chloroform/methanol mix (2:1; -20°C) were added to each sample in glass tubes and incubated on ice for 5 min. Then, samples were vortexed for 1 min and sonicated for 5 min prior to centrifugation (swinging, 2500 x *g*, 15 min). Resulting phases were collected separately using glass pipettes and stored in glass tubes for 10 min on ice. For the "polar metabolites" and "lipids" phases, the solvent was evaporated to obtain the crude extracts which were then solubilized at 10 $mg.mL^{-1}$ in MeOH/$H_2O$ 50:50 (*v/v*) or pure MeOH respectively, and then stored at -80°C until further analysis. For each experiment, a quality control sample (QC) was done by mixing 10 µL of every sample used in the experiment. The same protocol was applied without bacteria or EV samples in order to obtain extraction blanks for analysis. For protein phases, dried protein pellets were resuspended in 1 mL of cold pure MeOH (-20°C), vortexed and centrifugated (10 min, 12 000 x *g*). Supernatants were removed and remaining pellets were dried and then stored at -80°C until further analysis.

#### 3. Protein analyses

10 or 20 µL of protein samples were mixed with a Lysis buffer provided in the easyPep mini Kit (Thermo Scientific) to obtain a final volume of 100 µL. The samples were reduced and alkylated by incubation at 95°C for 10 min and continuously shaken at 1000 rpm. Then samples were digested with a LysC/Trypsin mixture enzyme for 3 h at 37°C and processed following the manufacturer's protocol. After a cleaning step, peptides were dried, suspended in 0.1% Formic Acid (FA) and dosed with the quantitative fluorometric peptide assay (ref. 23290, Thermo Scientific). 200 ng of each sample were analyzed on the Exploris 480 mass spectrometer coupled with a Vanquish NEO nanoLC system (Thermo Scientific). Peptides samples were loaded on a C18 Acclaim PepMap100 trap-column 300 µm ID x 5 mm, 5 µm,

100Å (ThermoFisher Scientific) and separated on a easyspray C18 Acclaim Pepmap100 nano-column, 50 cm x 75 µm i.d, 2 µm, 100 Å (Thermo Scientific) with a 22.5 minutes linear gradient from 3%  to 25% buffer B (A: 0.1% FA in $H_2O$, B: 0.1% FA in ACN/$H_2O$ (80/20)) from 25% to 35% of B in 7.5 min and then from 35% to 100% of B in 0.1 min, hold for 12 min and followed by a washing and equilibration steps for a total duration of 42 minutes. The flow rate was 300 $nL.min^{-1}$ and the oven temperature was kept constant at 45°C. Peptides were analysed with a DDA 1s HCD method: MS data were acquired in a data dependent strategy (DDA) selecting the fragmentation events based on the most abundant precursor ions in a 1s survey scan (350-1400 Th). Resolutions of the survey and MS/MS scans were respectively set at 120,000 and 15,000 at *m/z* 200 Th. The Ion Target Values for the survey and the MS/MS scans in the Orbitrap were set to 3E6 (300%) and 1E5 (100%) respectively and the maximum injection time was set to 50 ms for MS scan and 22 ms for MS/MS scan. Parameters for acquiring HCD MS/MS spectra were as follows: collision energy = 30 and isolation window = 4 *m/z*. The precursors with unknown charge state, charge state of 1 and 6 or greater than 6 were excluded. Peptides selected for MS/MS acquisition were then placed on an exclusion list for 40 s using the dynamic exclusion mode to limit duplicate spectra. Raw data were processed with Proteome Discoverer 3.1 (Thermo Scientific) through the CHIMERYS 3.0 search engine against the *Agrobacterium fabrum* database (Uniprot, Release January 2025) and a database of common contaminants. Precursor mass tolerance was set at 10 ppm and fragment mass tolerance was set at 0.02 Da, and up to 1 missed cleavages were allowed. Oxidation (M) was set as variable modification and Carbamidomethylation (C) as fixed modification. Validation of identified peptides and proteins was done using a target decoy approach with a false discovery rate positive (FDR < 1%) via Percolator. Protein quantitation was performed with precursor ions quantifier node in Proteome Discoverer 3.1 software. Proteins were normalized to the total peptide amount and ratios of quantitation were calculated in a pairwise way. The Match Between Run (MBR) was not applied. Statistical validation was based on t-test and a protein is considered differentially expressed between two conditions if the fold change is > 2 or < 0.5 and has a $p < 0.05$. Protein localization was predicted using DeepLocPro[2].

#### 4. *A. fabrum* C58 polar and apolar metabolites analyses

Dried polar and apolar fractions were solubilized at 10 $mg.mL^{-1}$ in MeOH 50:50 (*w/v*) and MeOH 100 respectively. For each fraction, a quality control sample (QC) was done by mixing 10 µL of every sample used in the experiment. The analysis of *A. fabrum* C58 metabolites was performed by UHPLC-DAD-ESI-MS-QTOF on an UHPLC Agilent 1290 coupled to a UV-vis Diode Array Detector (DAD) (Agilent 1290 Infinity series) and a high-resolution Q-TOF 6546 mass spectrometer (Agilent Technologies, Santa Clara, CA, USA). Liquid chromatography was carried out using a Poroshell 120 EC-$C_{18}$ column (3×100 mm, particle size 2.7 µm, Agilent technologies), preceded by a Poroshell $C_{18}$ guard column (3×5 mm, particle size 2.7 µm, Agilent technologies) maintained at 40°C with an injection volume of 4 µL per sample. For the polar fractions, the mobile phase was a mixture of acidified acetonitrile ($CH_3CN$) and acidified $H_2O$ (0.1% formic acid for each) applied at a flow-rate of 0.8 $mL.min^{-1}$, and with a gradient (acidified $H_2O$: acidified $CH_3CN$, [*v/v*]) starting at 99:1 for 1.5 min, increasing to 82:18 in 9.5 min, ramping up to 0.100 in 7 min and maintained for 2 min before returning to the starting conditions and equilibrating for 2.5 min. For the apolar fractions, the mobile phase was a mixture of $CH_3CN/H_2O$ (40:60 [*v/v*], 10 mM ammonium acetate) and $CH_3CN$/isopropanol

(10:90 [*v/v*], 10 mM ammonium acetate) applied at a flow-rate of 0.8 mL.min$^{-1}$, and with a gradient ($CH_3CN/H_2O$: $CH_3CN$/isopropanol, [*v/v*]) starting at 65:35 for 2 min, increasing to 0:100 in 24 min and maintained for 2 min before returning to the starting conditions and equilibrating for 1.5 min. QC sample was initially analyzed and then injected after every 6 samples in the run sequence to monitor the repeatability of the analysis. The quadrupole time-of-flight mass spectrometer (QTOF-MS) equipped with an electrospray ionization source (Dual AJS ESI Agilent, Santa Clara, CA, USA) was used in positive ionization mode $MS^2$ analyses under the following conditions: drying gas ($N_2$) flow of 12 L.min$^{-1}$ at 320 °C, nebulizer pressure of 40 psi, sheath gas flow rate of 11 L.min$^{-1}$ at 350 °C, with the capillary, nozzle and fragmentor voltages set to 3500 V, 500 V, and 100 V, respectively. The acquisition mass range was from *m*/*z* 50 to 2,500 with a scan rate of 4 spectra.s$^{-1}$. The $MS^2$ experiment was done with the collision energy set at 30 V, the $MS^2$ scan rate of 4 spectra.s$^{-1}$ and max precursors per cycle set at 4. Reference solutions containing standard compounds (HP-0921, *m/z* 922.0098 and purine, *m/z* 121.0509) were constantly infused as an accurate mass reference. The UHPLC-DAD-ESI-MS-QTOF device was managed by the Agilent Mass Hunter DataAcquisition version 11.0 software.

#### 5. Extraction of nopaline from tomato tumors

Tumors were harvested before being quickly frozen in liquid nitrogen before being dried by lyophilization overnight then crushed with stainless steel grinding balls (FastPrep-24 5G, MP Biomedicals, USA). Tumor powder samples (30 mg) were subjected to two successive extractions by adding 1 mL of MeOH/$H_2O$ 20:80 (*v/v*) , with 15 min sonication at each extraction step. After centrifugation (10 min, 19,745 × *g*), the 2 supernatants obtained were pooled. The solvent was evaporated to obtain the crude extracts which were then solubilized at 10 mg.mL$^{-1}$ in MS-grade $H_2O$ and stored at -80°C until further analysis.

#### 6. Extraction of *S. lycopersicum* metabolites

Eighteen hours post-inoculation, the 60 plants of each condition were randomly gathered in 6 groups to reduce variability prior to roots and aerial parts harvest. Roots and aerial parts were harvested separately before being quickly frozen in liquid nitrogen for further analyses. The plant parts were then freeze-dried overnight and crushed with stainless steel grinding balls (FastPrep-24 5G, MP Biomedicals, USA). For each experiment, plant powder samples were subjected to successive extractions by adding twice 1 mL of acidified MeOH/$H_2O$ 80:20 (*v/v*) (0.1% formic acid), and then twice 1 mL of acidified MeOH 100% (0.1% formic acid) with 15 min sonication at each extraction step. After centrifugation (10 min, 19,745 × *g*), the 4 supernatants obtained were pooled and transferred into a 5 mL hemolyze tube. The solvent was evaporated to obtain the crude extracts which were then solubilized at 10 mg.mL$^{-1}$ in MeOH/$H_2O$ 80:20 (*v/v*) and stored at -80°C until further analysis. For each experiment, a QC was done by mixing 6 µL of every sample used in the experiment.

#### 7. Targeted analysis of free amino acid content of *S. lycopersicum*

Tomato plant free amino acids were analyzed as described previously[3]. The analysis was performed on an HPLC Agilent 1100 coupled to a DAD and a fluorometric detector (FLD) (Agilent 1100 Infinity series). Norvaline (2 µL, 2.5 mM) was used as an internal standard and was added to 100 µL of the tested crude extract sample. Briefly, liquid chromatography was carried out using a Zorbax Eclipse-AAA column (4.6×150 mm, particle size 3.5 µm, Agilent technologies), preceded by a Zorbax Eclipse-AAA guard column (4.6x12.5mm, particle size 5µm, Agilent technologies) maintained at 40°C with an injection volume of 0.5µL per sample. A mix of 24 amino acids (α-aminobutyric acid, alanine, arginine, asparagine, aspartic acid, citruline, γ-aminobutyric acid, glutamine, glutamic acid, glycine, histidine, isoleucine, leucine, lysine, methionine, hydroxyproline, ornithine, phenylalanine, proline, serine, threonine, tryptophan, tyrosine, valine) prepared as described previously (**Henderson et *al.*, 2000**) was injected as standard mix. The HPLC-UV/DAD-FLD device was managed by Agilent ChemStation LC Rev. B.04.03-SP2 software, and the data was converted and reworked with MassHunter Qualitative Analysis B.07.00 software (Agilent Technologies).

#### 8. Analysis of specialized metabolites content of *S. lycopersicum*

The analysis of tomato plant specialized metabolites was performed by UHPLC-ESI-MS/MS-QTOF on an UHPLC Agilent 1290 coupled to a DAD (Agilent 1290 Infinity series) and a high-resolution Q-TOF 6546 mass spectrometer (Agilent Technologies, Santa Clara, CA, USA). Liquid chromatography was carried out using a Poroshell 120 EC-$C_{18}$ column (3×150 mm, particle size 2.7 µm, Agilent technologies), preceded by a Poroshell $C_{18}$ guard column (3×5 mm, particle size 2.7 µm, Agilent technologies) maintained at 45°C with an injection volume of 3 µL per sample. For tomato root metabolites, the mobile phase was a mixture of acidified $CH_3CN$ and acidified $H_2O$ (0.1% formic acid for each) applied at a flow-rate of 0.8 mL.min$^{-1}$, and with a gradient (acidified $H_2O$: acidified $CH_3CN$, [*v/v*]) starting at 99:1 for 1.5 min, increasing to 90:10 in 1.5 min, going to 84:16 in 7 min, going to 82:18 in 2.5 min, going to 73:27 in 7.5 min and maintained for 2 min, then going to 68:32 in 1 min, going to 0:100 in 3 min and maintained for 2 min, before returning to the starting conditions and equilibrating for 2 min. For tomato aerial part metabolites, the same solvents were applied at a flow-rate of 0.8 mL.min$^{-1}$ with a gradient (acidified $H_2O$: acidified $CH_3CN$, [*v/v*]) starting at 99:1 for 1.5 min, increasing to 93:7 in 1.5 min, going to 84:16 in 7 min, going to 82:18 in 2.5 min, going to 73:27 in 7.5 min and maintained for 2 min, then going to 68:32 in 1 min, going to 0:100 in 3 min and maintained for 2 min, before returning to the starting conditions and equilibrating for 2 min. For each experiment, the QC sample was initially analyzed and then injected after every 6 samples in the run sequence to monitor the repeatability of the analysis. The quadrupole time-of-flight mass spectrometer (QTOF-MS) equipped with an electrospray ionization source (Dual AJS ESI Agilent, Santa Clara, CA, USA) was used in positive ionization mode $MS^2$ analyses under the following conditions: drying gas ($N_2$) flow of 12 L.min$^{-1}$ at 320 °C, nebulizer pressure of 40 psi, sheath gas flow rate of 11 L.min$^{-1}$ at 350 °C, with the capillary, nozzle and fragmentor voltages set to 3500 V, 500 V, and 100 V, respectively. The acquisition mass range was *m*/*z* 50 to 2,500 with a scan rate of 4 spectra.s$^{-1}$. The $MS^2$ experiment was done with the collision energy set at 30 V, the $MS^2$ scan rate of 4 spectra.s$^{-1}$ and max precursors per cycle set at 4. Reference solutions containing standard compounds (HP-0921, *m*/*z* 922.0098 and purine, *m/z* 121.0509) were constantly infused as an accurate mass reference. The UHPLC-ESI-MS-

QTOF device was managed by the Agilent MassHunter Data Acquisition version 11.0 software. Data was processed using MzMine v4.5.0 for specialized metabolites profiles matrix obtention[4].

#### 9. Metabolite annotation and molecular networking

MS² data files were converted from the Agilent .d format to .mzML format using MSconvert software from the ProteoWizard package[5]. All .mzML files were processed using MzMine 4 v4.5.0[4]. The mass detection threshold was set to 1000 for MS and 50 for MS². The ADAP chromatogram builder was used with a minimum group size of scans of 3, a group intensity threshold and a minimum highest intensity of 10 000, and an *m/z* tolerance of 15 ppm. MS² scans pairing was done using a *m/z* tolerance of 15 ppm and a RT tolerance range of 0.2 min. Isotopes were filtered using the 13C isotope filter with an *m/z* tolerance of 25 ppm and a RT tolerance of 0.1 min. Peak alignment was performed using the join aligner module with a *m/z* tolerance of 25 ppm, weight for *m/z* of 80, RT tolerance of 0.1min and a weight for RT of 20. Features from the blanks were subtracted with a minimum detection in blanks of 1 occurrence and a fold-change threshold of 10. The resulting feature list was gap-filled with an intensity tolerance of 0.15, an *m/z* tolerance of 15 ppm, a RT tolerance of 0.1 min and a minimum scan of 1. Gap-filled feature list was then filtered to keep only features present in at least 6 samples and remaining 13C isotopes were also removed. The metaCorrelate and Ion identity networking algorithms were used with default parameters to assign adducts and correlation groups to the final feature list. The resulting .mgf and .csv metadata files were exported using the Export molecular networking file module. Sirius 6.1 was used to process the .mgf file and features of interest were annotated using the Molecular Formula Identification (Instrument Q-TOF, MS2 mass accuracy 10 ppm), ZODIAC, CANOPUS and CSI:FingerID modules[6–9]. Automatic annotations were manually verified using Lotus, Knapsack, GNPS, CasSciFinder databases, literature and analyses of standards when available. The .mgf and .csv files were imported into the MetGem v1.5.2 software[10]. MS² spectra were window-filtered by choosing only the top six peaks within the ± 50 Da window throughout the spectrum. The data were filtered by removing all peaks in the ± 17 Da range around the precursor *m/z*. The *m/z* tolerance window used to find the matching peaks was set to 0.05 Da, and cosine scores were kept under consideration for spectra sharing 3 matching peaks at least. The t-SNE molecular network was constructed by keeping nodes sharing at least 3 cosine scores above 0.5 with others. The number of iterations, perplexity, learning rate and early exaggeration parameters were set to 10 000, 6, 200 and 12 respectively. The Barnes-Hut approximation was activated using an angle of 0.5°.

#### 10. Statistical analysis

For EV content analyses, proteins or metabolites were kept only if present in the 3 samples of at least one condition at a minimal abundance of $10^7$ (proteins) or $10^3$ (metabolites). Normality was tested using Shapiro-Wilk's test. Subsequently, four tables representing the four tested comparisons (WCLGlu-WCLVir, EVGlu-EVVir, EVGlu-WCLGlu and EVVir-WCLVir) were created. The difference of abundance between two conditions was tested using Student's t-test or Wilcoxon test and Log2(FC) were calculated. Molecules were considered statistically enriched in one condition compared to the other when $p < 0.05$ and Log2(FC) > 1. Volcano plots were created using the package EnhancedVolcano v1.18.0. Protein GO terms, protein transmembrane helices presence, KEGG levels, metabolic classes and metabolic synthesis pathway enrichment were made using a Fisher's test on differently accumulated proteins or metabolites for each comparison. Categories were considered enriched when $p < 0.05$. For tumor formation analyses, data were analyzed using a Linear Mixed-Effects Model (LMM) using the lme4 v1.1.37 and lmerTest v3.1 packages. The treatment solution was defined as a fixed effect, while the experimental week was included as a random effect to account for environmental variation between blocks. Each of the 27 plants was treated as an independent biological replicate. Post-hoc pairwise comparisons were performed using Tukey's HSD test via the emmeans v1.11.2 package. For the plant metabolites analyses, ions representative of metabolites were kept only if present in the 6 samples of at least one condition with a minimal abundance of $10^3$. Abundance values were Log10-transformed and pareto-scaled to create four tables corresponding to the four tested comparisons (Control-C58Glu-C58Vir, Control-EVGlu-EVVir, Control-EVGlu-C58Glu, Control-EVVir-C58Vir). Partial Least Squares Discriminant Analysis (PLS-DA) method from the R package mixOmics was used to identify discriminating variables between the tested conditions (VIP score > 1)[11]. For the annotated variables with a VIP score > 1, the difference of abundance was tested using ANOVA (Tukey's post-hoc test) or Kruskal-Wallis (Dunn's post-hoc test) test after assessing the normality using Shapiro-Wilks test.
